# Mechanism of Lys6 poly-ubiquitin specificity by the *L. pneumophila* deubiquitinase LotA

**DOI:** 10.1101/2022.05.11.491541

**Authors:** Gus D. Warren, Tomoe Kitao, Tyler G. Franklin, Justine V. Nguyen, Paul P. Geurink, Tomoko Kubori, Hiroki Nagai, Jonathan N. Pruneda

**Affiliations:** Department of Molecular Microbiology & Immunology, Oregon Health & Science University, Portland, OR 97239, USA; Department of Microbiology, Graduate School of Medicine, Gifu University, Gifu, Gifu 501-1194, Japan; Oncode Institute & Department of Cell and Chemical Biology, Leiden University Medical Centre, Leiden, The Netherlands; G-CHAIN, Gifu University, Gifu, Gifu 501-1194, Japan

**Author notes:** These authors contributed equally (Tomoe Kitao, Tyler G. Franklin). Address correspondence to Jonathan N. Pruneda.

**Keywords:** Ubiquitin, Deubiquitinase, Legionella pneumophila, Bacterial effector

## Abstract

The versatility of ubiquitination to impose control over vast domains of eukaryotic biology is due, in part, to diversification through differently-linked poly-ubiquitin chains. Deciphering the signaling roles for some poly-ubiquitin chain types, including those linked via K6, has been stymied by a lack of stringent linkage specificity among the implicated regulatory proteins. Forged through strong evolutionary pressures, pathogenic bacteria have evolved intricate mechanisms to regulate host ubiquitin, and in some cases even with exquisite specificity for distinct poly-ubiquitin signals. Herein, we identify and characterize a deubiquitinase domain of the secreted effector protein LotA from *Legionella pneumophila* that specifically regulates K6-linked poly-ubiquitin during infection. We demonstrate the utility of LotA as a tool for studying K6 poly-ubiquitin. By determining apo and diUb-bound structures, we identify the mechanism of LotA activation and K6 poly-ubiquitin specificity, and identify a novel ubiquitin-binding domain utilized among bacterial deubiquitinases.

## INTRODUCTION

Post-translational regulation with the small protein modifier ubiquitin (Ub) is essential to all eukaryotic life. Breakdown of this process can lead to many forms of severe disease, including cancer, neurodegeneration, and autoimmunity (Popovic *et al*, 2014). In humans, many hundreds of enzymes tightly regulate the process of protein ubiquitination by defining not only the target substrate but also the type of Ub modification (Clague *et al*, 2015). Unlike binary “on-off” modifications such as phosphorylation or acetylation, the Ub signal can be further diversified through the formation of polymeric Ub (polyUb) chains in which any of the seven Lys residues (e.g., K48- or K63-linked chains) or the amino-terminus (linear/M1-linked chains) are ubiquitinated. Differently-linked polyUb chains code for unique signaling outcomes, including proteasomal degradation (K48-linked chains), the DNA damage response (K63-linked chains), or innate immune signaling (M1-linked chains) (Komander & Rape, 2012). Although the existence and importance of all eight possible chain types has been established (Xu *et al*, 2009; Wagner *et al*, 2011; Roscoe *et al*, 2013; Meza Gutierrez *et al*, 2018), signaling outcomes and regulators for half (K6, K27, K29, and K33) remain ill-defined and have thus been coined “atypical” chains (Swatek & Komander, 2016).

Among the atypical chain types, K6-linked polyUb has remained particularly enigmatic. Whether K6 polyUb serves as a proteasomal degradation signal remains unclear, as levels of K6 chains show little or no increase upon proteasomal inhibition (Kim *et al*, 2011; Wagner *et al*, 2011). K6 polyUb has been shown to accumulate following mitochondrial depolarization, UV-induced DNA damage, or inhibition of the AAA ATPase p97 (Ordureau *et al*, 2014; Elia *et al*, 2015; Heidelberger *et al*, 2018; Michel *et al*, 2017). Exact description of the importance of K6 polyUb in these responses, however, has been clouded by the imprecise specificities of the E3 ligases and deubiquitinases (DUBs) proposed to be involved. Mitophagy, for example, can be regulated by the E3 ligase Parkin and DUBs USP30 and USP8, and while these enzymes can control levels of K6 polyUb, they also target other chain types including K11, K48, and K63 (Ordureau *et al*, 2014; Bingol *et al*, 2014; Cunningham *et al*, 2015; Gersch *et al*, 2017; Durcan *et al*, 2014). The same is true for the ligase activity of BRCA1 in the DNA damage response (Wu-Baer *et al*, 2003; Morris & Solomon, 2004; Nishikawa *et al*, 2004), as well as the ligase activity of HUWE1 in targeting substrates for p97 (Heidelberger *et al*, 2018; Michel *et al*, 2017; Jäckl *et al*, 2018). Ascribing a signaling function for K6 polyUb with high confidence, therefore, awaits the discovery of new approaches or enzymes that can more specifically measure and/or perturb it. Such was the case with M1-linked polyUb, prior to the discovery of M1-specific regulators (Elliott, 2016).

PolyUb chain specificity is typically achieved through the concerted action of multiple Ub-binding sites. Within the context of a polyUb chain, Ub molecules on either side of an (iso)peptide linkage can be classified as a distal Ub (Ub^dist^) that is linked via its C-terminus onto an amino group of the proximal Ub (Ub^prox^). Both E2 Ub-conjugating enzymes and E3 ligases achieve linkage specificity through an acceptor Ub-binding site that orients what will become Ub^prox^ for nucleophilic attack on what will become Ub^dist^, which is bound into the donor site via a high-energy E2/E3∼Ub thioester linkage (Deol *et al*, 2019). For DUBs, specificity is often achieved via Ub-binding sites on opposite sides of the catalytic center, including an S1 site that binds Ub^dist^ and an S1’ site that binds Ub^prox^ in a linkage-specific orientation (Mevissen & Komander, 2017). Linkage-specific regulators have been identified for M1, K11, K48, and K63 polyUb (Komander & Rape, 2012), but despite considerable efforts, enzymes with stringent specificity toward K6 polyUb in humans have thus far evaded detection.

Pathogenic bacteria may represent an alternative source of polyUb linkage-specific enzymes. Bacterial pathogens have evolved Ub regulatory enzymes that interfere with human signaling processes during infection, including a number that target specific polyUb chain types (Franklin & Pruneda, 2021). In fact, one such enzyme from enterohemorrhagic *Escherichia coli* called NleL is the most K6-specific ligase known to-date, assembling a 50:50 mixture of K6 and K48 linkages (Lin *et al*, 2011). The causative agent of Legionnaires’ disease, *Legionella pneumophila*, has evolved numerous mechanisms of disrupting or hijacking host Ub signaling. Of the >300 effector proteins secreted into infected cells via the *L. pneumophila* Dot/Icm type IV secretion system, several dozen have been shown to regulate host Ub signals (Qiu & Luo, 2017), including some that target specific polyUb chain types (Wan *et al*, 2019). Many of these Ub-targeted *Legionella* effectors have been shown to localize to the *Legionella*-containing vacuole (LCV), where they edit the complex “coat” of Ub modifications in order to establish a replicative niche and evade host immune responses (Kitao *et al*, 2020). One such effector, termed *<u>L</u>egionella* <u>OT</u>U-like protein <u>A</u> (LotA), was reported to contain two catalytic DUB domains with similarity to the human OTU family, as well as activity toward K6, K48, and K63 polyUb (Kubori *et al*, 2018). Whether and how one of LotA’s catalytic OTU domains might be specific toward K6 polyUb, however, remained unknown.

Here we demonstrate that the two catalytic DUB modules of LotA are separable and have unique activities and specificities. Remarkably, the first OTU domain of LotA encodes perfect specificity toward K6 polyUb and represents a valuable tool for studying this atypical linkage. To understand the molecular basis for K6 specificity, we determined crystal structures for LotA alone and bound to K6 diUb. Detailed biochemical analyses revealed a multi-step mechanism of LotA activation that endows K6 specificity, which incorporates themes of conformational rearrangement and substrate-assisted catalysis observed in human linkage-specific DUBs. In the process of studying LotA, we discovered a new class of Ub-binding domain that is adapted across the *Legionella* Lot class as well as an otherwise unrelated DUB from *Orientia tsutsugamushi*. This work provides a powerful new tool for the study of Ub biology and firmly establishes an interesting link for K6 polyUb at the host-pathogen interface.

## RESULTS

### LotA encodes two catalytic DUB domains with distinct specificities

The Lot class of DUBs encoded by *L. pneumophila* is currently composed of LotA (lpg2248), LotB (lpg1621), LotC (lpg2529), and LotD (lpg0227) and defined by a ∼100 amino acid insertion within the OTU domain. The Lot-class insertion domains are more diverse in primary sequence than the core OTU domains (**Fig. S1A-B**), suggesting that they may play a role in determining the unique polyUb specificities that have been reported (Kubori *et al*, 2018; Ma *et al*, 2020; Schubert *et al*, 2020; Shin *et al*, 2020; Liu *et al*, 2020; Hermanns *et al*, 2020). LotA encodes two Lot-class OTU domains, denoted LotA_N_ and LotA_M_ for the N-terminal and middle domains, respectively, followed by the C-terminal LotA_C_ domain that binds PI(3)P, localizing LotA to the LCV during infection (Kubori *et al*, 2018) (**Fig. 1A**). The sequence relationship between LotA_N_ and LotA_M_ is the most distinct observed among the Lot class, with sequence identities of only 19% and 23% within the insertion and OTU domains, respectively (**Fig. S1A-B**). To determine if LotA_N_ and LotA_M_ contribute distinct activities to the K6-, K48-, and K63-targeted DUB activity reported for the full length LotA (Kubori *et al*, 2018), we cloned and expressed the isolated domains for biochemical analyses.

**Figure 1:**
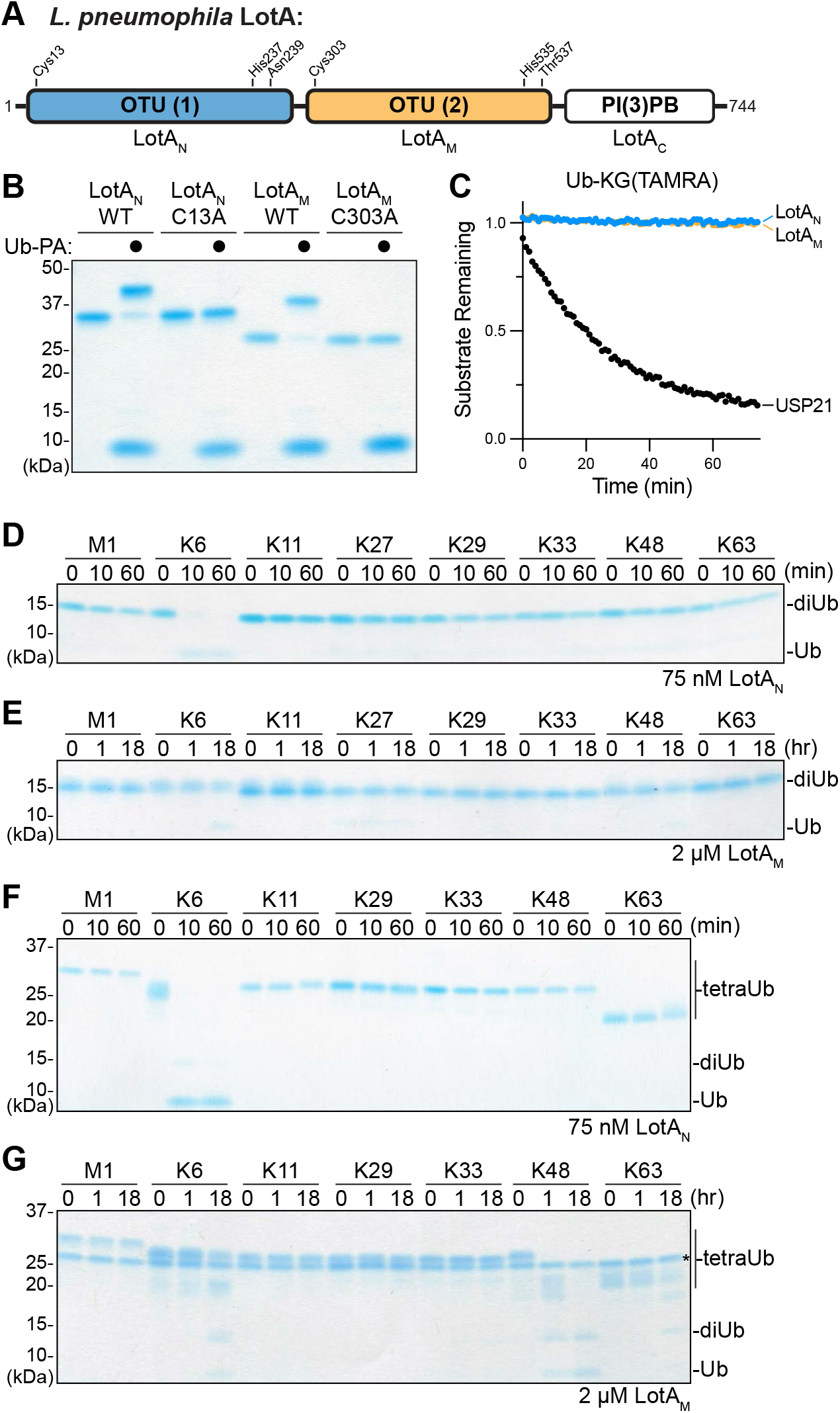
Separation of LotA deubiquitinase activities. A. Domain architecture of *L. pneuomophila* LotA, with catalytic triad residues annotated for each of the two OTU domains. B. Coomassie-stained SDS-PAGE gel showing purified constructs of LotA_N_ (1-300) and LotA_M_ (287-543), alongside their catalytically-inactive variants. Reactivity with the Ub-PA activity-based probe is assessed by a shift in mobility. C. Ub-KG(TAMRA) cleavage assay monitored by fluorescence polarization. Activity was measured using 1 µM LotA_N_, 1 µM LotA_M_, or 100 nM USP21 as a control DUB. D-E. Gel-based polyUb chain specificity analysis against all eight canonical diUb linkages. Reactions containing the indicated concentrations of LotA_N_ (**D**) and LotA_M_ (**E**) were sampled at the indicated timepoints, quenched, and resolved by SDS-PAGE with Coomassie staining. F-G. Gel-based polyUb chain specificity analysis against seven tetraUb linkages. Reactions containing the indicated concentrations of LotA_N_ (**F**) and LotA_M_ (**G**) were sampled at the indicated timepoints, quenched, and resolved by SDS-PAGE with Coomassie staining. See also Figure S1.

Both the LotA_N_ and LotA_M_ isolated constructs reacted with the Ub-Propargylamide (Ub-PA) activity-based probe, indicating that each are competent for DUB activity (**Fig. 1B**) (Ekkebus *et al*, 2013). Reactivities of LotA_N_ and LotA_M_ with Ub-PA were ablated by mutation of their annotated catalytic cysteine residues, C13 and C303, to alanine (**Fig. 1A-B**) (Kubori *et al*, 2018). Surprisingly, although both LotA_N_ and LotA_M_ were able to bind and react with the monomeric Ub-PA substrate, neither could cleave the monomeric Ub-KG(TAMRA) fluorescent substrate, or the analogous NEDD8, SUMO1, or ISG15 substrates (a discrepancy that will be discussed further below) (**Fig. 1C, S1C-E**) (Geurink *et al*, 2012). When incubated with each of the eight canonical (iso)peptide-linked diUb substrates, LotA_N_ demonstrated remarkable specificity for the K6 linkage type (**Fig. 1D**). In contrast, against the same panel of diUb substrates, LotA_M_ was very inefficient and only showed marginal cleavage of the K6- and K48-linked substrates at high concentration after a long incubation period (**Fig. 1E**). As the polyUb chain length has been reported to affect the activities and/or specificities of some DUBs (Mevissen *et al*, 2013; Békés *et al*, 2015; Abdul Rehman *et al*, 2016), we assembled a panel of seven differently-linked tetraUb substrates (lacking only K27). LotA_N_ again showed exquisite specificity against the K6 linkage, with no detectable activity toward any of the other substrates (**Fig. 1F**). Against the longer chain length, LotA_M_ now demonstrated more robust DUB activity, with a distinct preference toward the K48 linkage and lesser activities toward K6 and K63 tetraUb (**Fig. 1G**). Consistent with observations made with the diUb panel (**Fig. 1E**), LotA_M_ activity appeared to stall after the tetraUb and triUb were consumed, leaving a diUb product that was stable over time (**Fig. 1G**). These results indicate that LotA_N_ and LotA_M_ indeed demonstrate unique DUB properties, with LotA_N_ showing exquisite K6 specificity while LotA_M_ shows a preference toward longer K48-linked polyUb that is likely directed by an additional (S2 or S2’) Ub-binding site.

### Benchmarking LotA_N_ as a tool for the study of K6 polyUb

To further characterize the activity of LotA_N_ toward K6 polyUb, we constructed a fluorescent K6 diUb substrate (see Materials and Methods) and monitored cleavage by fluorescence polarization (FP), as reported previously (Virdee *et al*, 2010; Keusekotten *et al*, 2013; Pruneda *et al*, 2016). Through measuring initial rates of catalysis over a range of substrate concentrations, Michaelis-Menten parameters could be extracted for LotA_N_ against K6 diUb (**Fig. 2A, S2A**). Despite a relatively high K_M_, LotA_N_ demonstrated an overall catalytic efficiency that is similar to other linkage-specific OTU DUBs, including the K11-specific Cezanne and the M1-specific OTULIN (Mevissen *et al*, 2016; Keusekotten *et al*, 2013).

**Figure 2:**
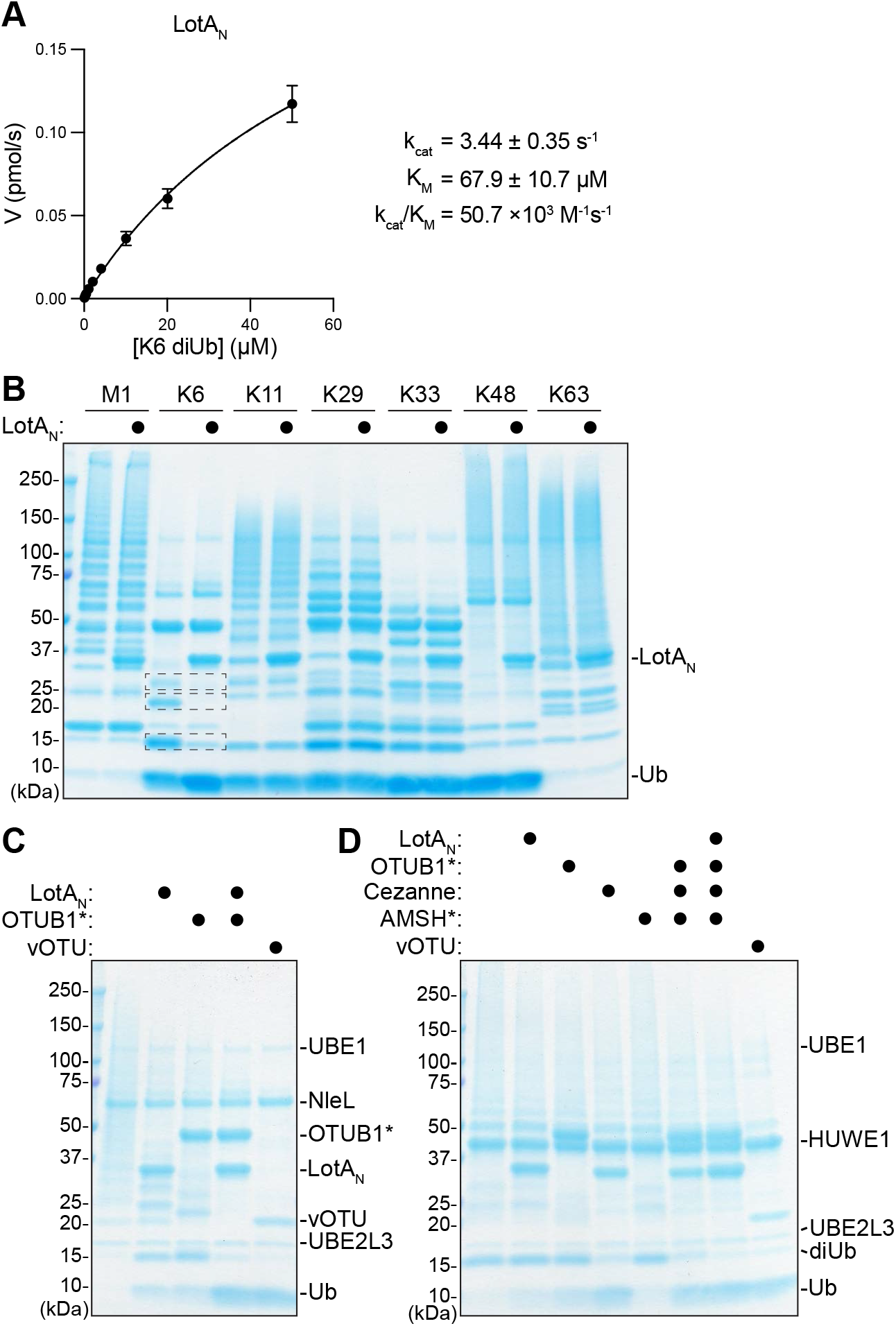
Application of LotA_N_ K6 specificity for UbiCRest analysis. A. Kinetic parameters of LotA_N_ (1-300) measured by changes in fluorescence polarization of labeled K6 diUb. Initial rates of diUb cleavage were measured over a range of LotA_N_ concentrations and fit to a Michaelis-Menten model. Error bars represent standard deviation over series of measurements, each made in triplicate. B. Homogeneous assemblies for seven polyUb linkage types were generated and treated with high concentration (5 µM) LotA_N_ for 2 h before the reactions were quenched, resolved by SDS PAGE, and visualized for cleavage by Coomassie staining. In the LotA_N_ treatment of K6 polyUb, all observed polyUb bands are indicated with boxes. All LotA_N_-resistant protein bands in the K6 polyUb treatment are enzymes used in the substrate assembly process. C. UbiCRest analysis of an NleL ligase assembly with 5 µM K6-specific LotA_N_, K48-specific OTUB1*, and nonspecific vOTU. Cleavage of NleL-assembled polyUb can be observed by a decrease in the produce “smear” or by a reappearance of monoUb. D. UbiCRest analysis of a HUWE1 ligase assembly with 5 µM K6-specific LotA_N_, K48-specific OTUB1*, K11-specific Cezanne, K63-specific AMSH*, and nonspecific vOTU. Combinations of all linkage-specific DUBs with and without LotA_N_ are also shown. Cleavage of HUWE1-assembled polyUb can be observed by a decrease in the produce “smear” or by a reappearance of monoUb. See also Figure S2.

Our specificity analysis of LotA_N_ against panels of diUb and tetraUb substrates showed remarkable preference of K6 linkages at low LotA_N_ concentrations. To further understand the specificity of LotA_N_ and benchmark its utility as a tool, we tested if specificity is retained at high enzyme concentrations against more complex polyUb substrates. Using established E2/E3/DUB combinations (Michel *et al*, 2018), we generated complex but homogeneous mixtures of polyUb chains for seven linkage types (excluding K27) and incubated them with a high concentration of LotA_N_ (**Fig. 2B**). Under these conditions, LotA_N_ still demonstrated exclusive cleavage of K6 linkages, demonstrating a high level of specificity even at elevated concentration.

To test LotA_N_’s utility in releasing K6 linkages from complex, heterogeneous mixtures, we tested it alongside other linkage-specific DUBs in several Ubiquitin Chain Restriction (UbiCRest) analyses (Hospenthal *et al*, 2013; Mevissen *et al*, 2013; Hospenthal *et al*, 2015). The HECT-like E3 ligase NleL from enterohemorrhagic *Escherichia coli* has been reported to assemble mixed K6 and K48 polyUb chains (Lin *et al*, 2011; Hospenthal *et al*, 2013). Alongside the constitutively-activated K48-specific DUB OTUB1* (Michel *et al*, 2015), LotA_N_ could indeed confirm this chain architecture in a UbiCRest assay. Treatment of NleL-assembled polyUb with either LotA_N_ or OTUB1* resulted in a partial collapse of the polyUb smear down to shorter chains such as diUb, as well as a partial return of the monoUb band (**Fig. 2C**). Co-incubation with both LotA_N_ and OTUB1* cleaved the vast majority the input polyUb, producing similar levels of released monoUb as treatment with the nonspecific DUB vOTU (Akutsu *et al*, 2011; Mevissen *et al*, 2013) (**Fig. 2C**).

To further test the utility of LotA_N_ in diagnosing the presence of K6 linkages among a complex mixture, we generated polyUb chains with HUWE1, a human HECT E3 ligase that is reported to assemble a mixture of K6-, K11-, and K48-linked chains (Michel *et al*, 2017; Jäckl *et al*, 2018; Grabarczyk *et al*, 2021). Treatment with the K11-specific DUB Cezanne released the largest amount of monoUb, followed by the K48-specific OTUB1*, then LotA_N_, and finally the constitutively-activated K63-specific AMSH* (Michel *et al*, 2015), which had only a marginal amount of activity (**Fig. 2D**). Co-incubation of OTUB1*, Cezanne, and AMSH* with or without LotA_N_ showed a slight difference attributable to K6 linkages, especially at the level of diUb products released by the other chain-specific DUBs (**Fig. 2D**). Treatment with vOTU does, however, appear to cleave more of the HUWE1-assembled products than the combined treatment of LotA_N_, OTUB1*, Cezanne, and AMSH*, suggesting either the presence of additional linkage types or mono-ubiquitinated substrates that cannot be processed by the chain-specific DUBs (**Fig. 2D**). Thus, the high level of activity and specificity exhibited by LotA_N_ makes it a powerful tool that can be used alongside other linkage-specific enzymes to study K6 polyUb.

### Structural analysis of LotA_N_

The unique sequence (**Fig. S1A-B**) and catalytic properties of LotA_N_ warranted structural analysis for comparison to the other Lot DUBs. We determined a 1.5Å crystal structure of LotA_N_ residues 1-294 using SAD phasing on bound iodide ions (**Fig. 3A, S3A, Table S1**). Residues 2-275 could be confidently modeled into the electron density. Overall, the LotA_N_ structure resembled other Lot DUBs, with a papain-like OTU domain and a characteristic α-helical insertion domain at Variable Region 1 (VR-1), but additionally revealed several unique features (**Fig. 3A**). Firstly, the catalytic triad, comprised of C13, H237, and N239, was misaligned and beyond hydrogen bonding distance in the structure. In fact, two conformations of the C13 side chain were visible in the electron density, whereas no electron density was observed for the H237 imidazole ring (**Fig. 3B**). Despite this, mutation of C13, H237, and to a lesser extent N239 all severely decreased the ability of LotA_N_ to cleave the fluorescent K6 diUb substrate (**Fig. 3C**). The catalytic triad misalignment observed in apo LotA_N_ combined with its inability to cleave the monomeric Ub-KG(TAMRA) substrate (**Fig. 1C**) is consistent with the possibility that binding of K6 diUb is required for rearrangement and activation of the catalytic center, a mechanism termed “substrate-assisted catalysis” that has been observed previously for the M1-specific OTULIN (Keusekotten *et al*, 2013).

**Figure 3:**
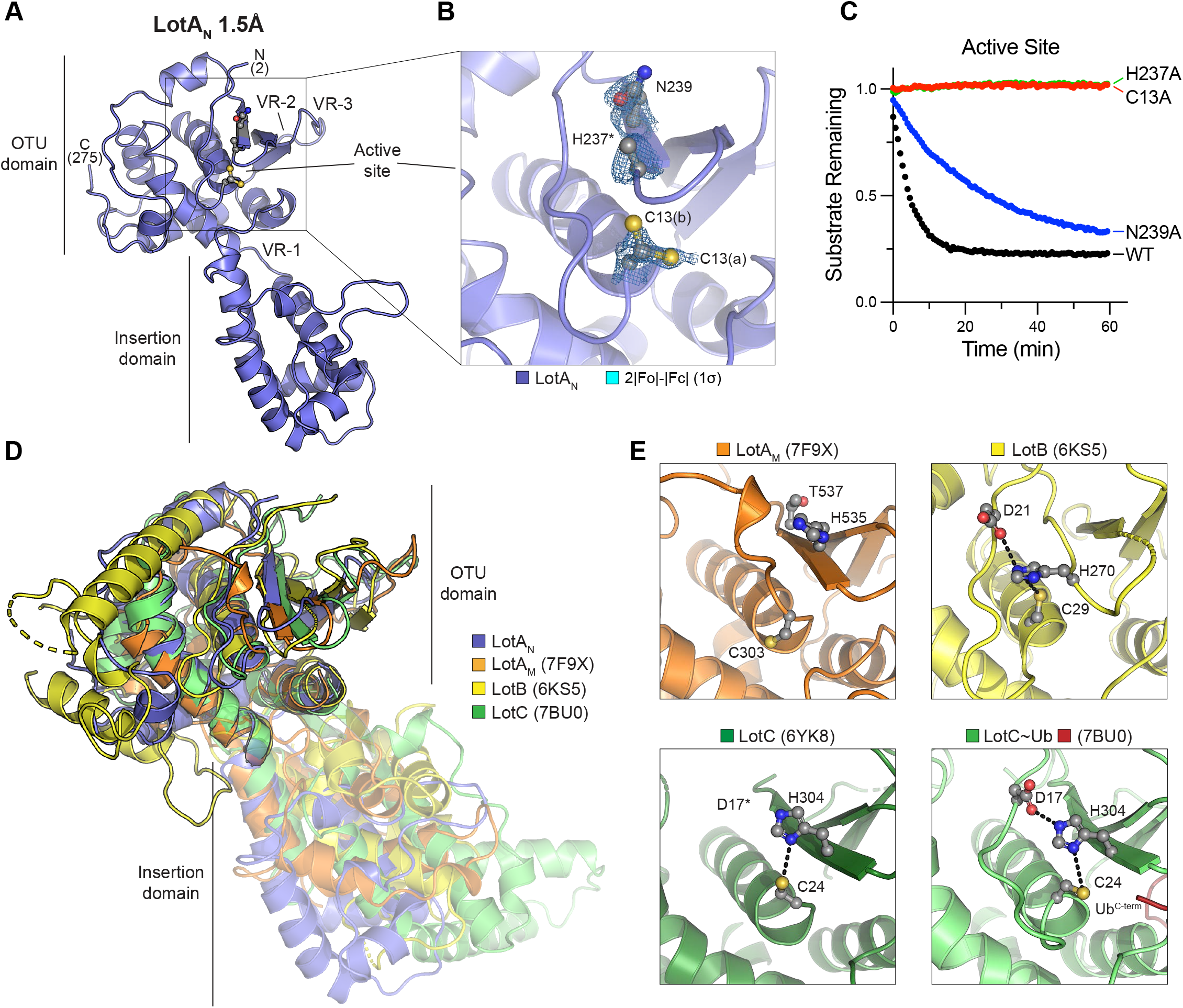
Crystal structure of the LotA_N_ OTU domain. A. 1.5Å crystal structure of LotA (1-294) determined by SAD, labeled with visible termini, domain architecture, active site, and OTU variable regions (VR1-3) described previously (Schubert *et al*, 2020). B. Close-up view of the LotA_N_ active site with 2|Fo|-|Fc| electron density overlaid at 1χρ for catalytic triad residues. Alternate conformations of the C13 side chain (a and b) are observed, whereas density for the H237 imidazole ring is absent. C. Cleavage of fluorescent K6 diUb by the indicated LotA_N_ active site variants at 10 nM concentration monitored by fluorescence polarization. D. Structural overlay of LotA_N_ (blue), LotA_M_ (PDB 7F9X, orange), LotB (PDB 6KS5, yellow), and LotC (PDB 7BU0, green). Structures were aligned by their core OTU domains to demonstrate variability in the insertion domains. E. Close-up views of the LotA_M_, LotB, LotC (PDB 6YK8), and Ub-bound LotC (PDB 7BU0) active sites. Catalytic triad residues are shown in ball-and-stick representation, with hydrogen bonds indicated by dashed lines. See also Figure S3.

The core OTU fold of LotA_N_ closely resembles those of the other Lot-class DUBs (**Fig. 3D**), as well as the most closely related OTU DUBs from humans, viruses, and bacteria outside of *Legionella* (Ma *et al*, 2020; Shin *et al*, 2020; Liu *et al*, 2020; Takekawa *et al*, 2022; Juang *et al*, 2012; Akutsu *et al*, 2011; Schubert *et al*, 2020; Holm, 2020) (**Fig. S3B-D**). At the putative S1 Ub-binding site, LotA_N_ more closely follows the other Lot-class DUBs with a large helical domain inserted at VR-1, and minimal structural features at VR-2 and VR-3 compared to other OTU domains (Schubert *et al*, 2020) (**Fig. 3D, S3B-D**). All of the Lot-class insertion domains structurally resolved thus far exhibit unique adaptions of an underlying helical sub-structure with distinct orientations with respect to the OTU domains (**Fig. 3D**), a feature that has been proposed to contribute to the unique polyUb specificities observed within this class of DUBs. A closer look at the catalytic triad across the related Lot-class DUBs reveals that, like LotA_N_, the active site of LotA_M_ is also misaligned in the observed crystal structure, whereas the active sites of LotB, LotC, and Ub-bound LotC are all in proper alignment for catalysis (**Fig. 3E**). This discrepancy in catalytic triad alignment is consistent with our observation that neither LotA_N_ nor LotA_M_ could cleave the monomeric Ub-KG(TAMRA) substrate (**Fig. 1C**), whereas LotB and the related LotD were previously shown to be active in the same assay (Schubert *et al*, 2020). This suggests that LotA_N_ and LotA_M_ are unique among the Lot-class DUBs in their requirement for some form of substrate-assisted catalysis.

### Mechanism of K6 diUb recognition by LotA_N_

To understand the molecular basis of LotA_N_ activation and K6 polyUb specificity, we determined a 2.8Å crystal structure of LotA_N_ residues 1-276 in complex with K6 diUb by molecular replacement (**Fig. 4A, S4A-B, Table S1**). In the structure, K6 diUb is bound across the LotA_N_ active site, sandwiched between the OTU core and the helical insertion domain (**Fig. 4A**). In order to preserve the K6 diUb while retaining binding to LotA_N_, the active site C13A mutation was used. Despite this, the electron density between the LotA_N_-bound Ub molecules more closely supports a structural intermediate just after isopeptide cleavage, with no density connecting the -amino group of K6 from Ub^prox^ with the carbonyl carbon of G76 in Ub^dist^ (**Fig. 4B**). Rather than cleavage of the K6 linkage, we believe this lack of clear electron density is either due to a shift in register (i.e., Ub^dist^ and Ub^prox^ are bound into the S1’ and S1 sites, respectively, of symmetry-related LotA_N_ molecules) or symmetry averaging across related asymmetric units, as the C-terminus of Ub^prox^ is in close proximity to the K6 of a symmetry-related Ub^dist^ (**Fig. S4C**). The symmetry-related expansion of polyUb chains has been observed in other crystal structures (Michel *et al*, 2015), and in this case suggests that LotA_N_ exhibits endo-DUB activity that can cleave anywhere within a polyUb chain. Although the catalytic C13 is mutated to alanine, the electron density clearly supports proper alignment and hydrogen bond distance of the remaining H237 and N239 triad members (**Fig. 4B**), indicating a structural rearrangement upon K6 diUb binding that will be discussed in more detail below.

**Figure 4:**
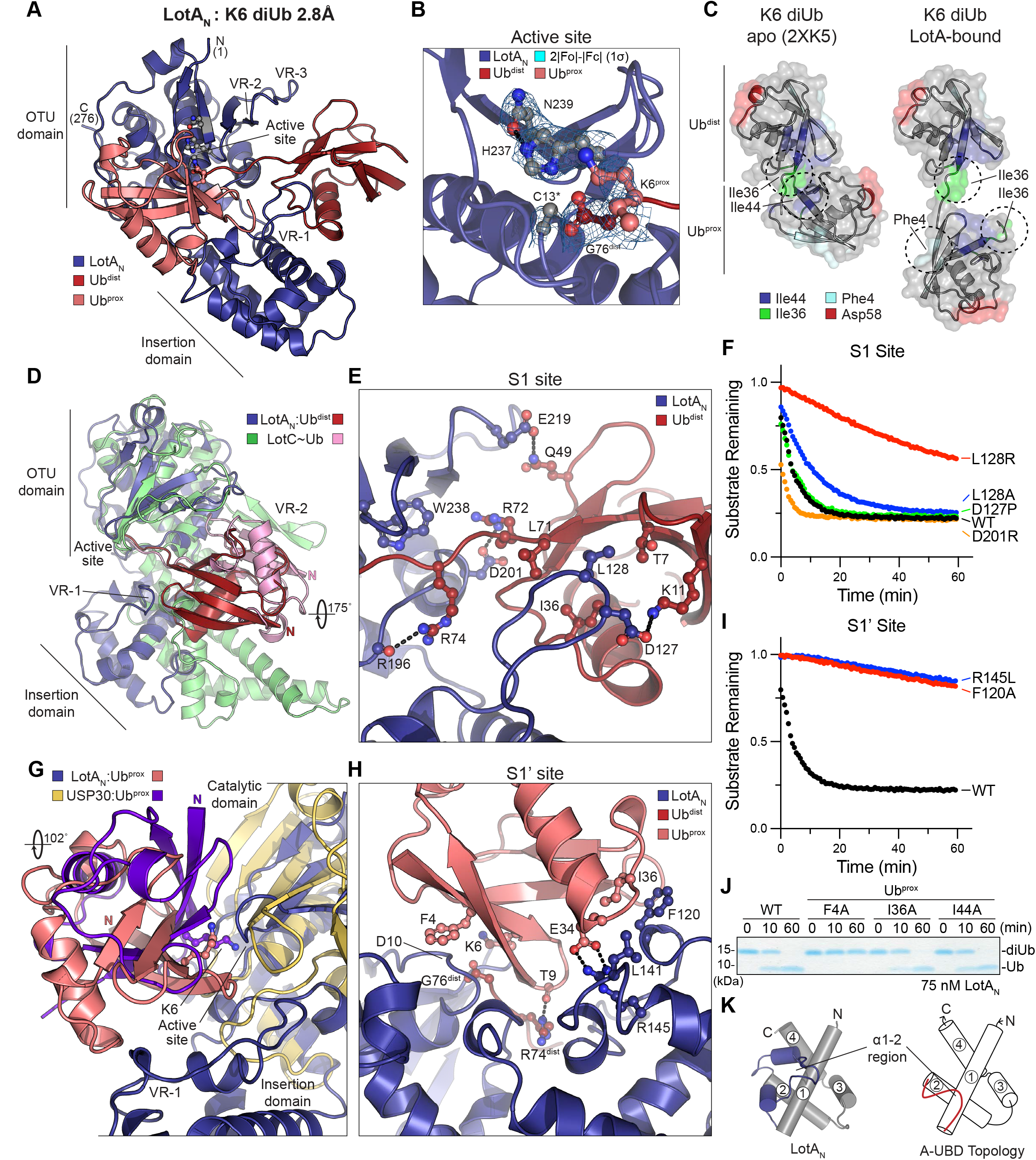
Crystal structure of the LotA_N_ OTU domain bound to K6 diUb. A. 2.8Å crystal structure of LotA_N_ (1-276, blue) bound to K6 diUb (shades of red), determined by MR. The structure is labeled with visible termini, domain architecture, active site, and OTU variable regions (VR1-3). B. Close-up view of the K6 diUb-bound LotA_N_ active site with 2|Fo|-|Fc| electron density overlaid at 1χρ for catalytic triad residues. The K6 side chain of Ub^prox^ and C-terminus of Ub^dist^ are also shown with overlaid electron density. The LotA_N_ catalytically inactive C13A variant was used to preserve the diUb linkage. C. Crystal structures of K6 diUb in isolation (PDB 2XK5) and bound to LotA_N_, aligned by their distal Ub moieties (top) and shown side-by-side. Availability and orientation of common interaction surfaces are shown, with dashed circles indicating the surfaces utilized by either Ub:Ub or LotA_N_:Ub interaction. D. Structural overlay of LotA_N_ (blue) and LotC (green) bound to their respective distal Ub moieties (red and pink, respectively). The structures are aligned on their core OTU domains, and highlight large differences in Ub orientation, insertion domains, and use of variable regions. E. Close-up view of the LotA_N_ S1 site (blue) bound to Ub^dist^ (red). Interacting residues are shown in ball-and-stick representation, with hydrogen bonds indicated by dashed lines. F. Cleavage of fluorescent K6 diUb by the indicated LotA_N_ S1 site variants at 10 nM concentration monitored by fluorescence polarization. G. Structural overlay of LotA_N_ (blue) and human USP30 (tan) bound to their respective distal Ub moieties (salmon and purple, respectively). The structures are aligned on their core catalytic domains, and highlight large differences in Ub orientation and features of their S1’ sites. H. Close-up view of the LotA_N_ S1’ site (blue) bound to Ub^prox^ (salmon). Interacting residues are shown in ball-and-stick representation, with hydrogen bonds indicated by dashed lines. I. Cleavage of fluorescent K6 diUb by the indicated LotA_N_ S1’ site variants at 10 nM concentration monitored by fluorescence polarization. These data were collected in parallel with those presented in (**F**), and the WT dataset is shown again for reference. J. Gel-based LotA_N_ cleavage assay of K6 diUb variants encoding the indicated Ub^prox^ mutations. Reactions were quenched at the indicated times and visualized by SDS PAGE with Coomassie staining. K. Underlying helical domain architecture of the LotA_N_ (left) and stereotypical (right) adaptive Ub-binding domain (A-UBD), with helices labeled and the Ub-binding α1-2 regions shown in color. See also Figure S4.

In isolation of binding partners, crystallography and solution NMR data support a predominately closed conformation of K6 diUb, in which the Ub^dist^ I36 hydrophobic patch is bound to the Ub^prox^ I44 hydrophobic patch (Virdee *et al*, 2010; Hospenthal *et al*, 2013) (**Fig. 4C**). In contrast, LotA_N_ appears to select an open conformation of K6 diUb such that it can interact with the Ub^dist^ I36 patch as well as the I36 and F4 hydrophobic patches of Ub^prox^ (**Fig. 4C**). Selection among heterogeneous K6 diUb conformations appears to be an emerging trend of K6-interacting proteins, as unique K6 diUb structures have been resolved in complexes with an engineered affimer protein, the DUB USP30, the NZF Ub-binding domain of TAB2, and now also LotA_N_ (Michel *et al*, 2017; Gersch *et al*, 2017; Li *et al*, 2021) (**Fig. S4D**). Each unique conformation of K6 diUb presents a unique configuration of Ub^dist^ and Ub^prox^ hydrophobic patches, and thus far the I44, I36, and F4 patches are the most commonly utilized for binding (**Fig. S4D**).

The interaction with Ub^dist^ at the LotA_N_ S1 site appears to be mainly driven through an interface between the inserted helical domain at VR-1 and the I36 patch of Ub^dist^ (**Fig. 4A, C**). In contrast, the structure of LotC bound to Ub shows a stark ∼180-degree rotation and a primary interface at a unique VR-2 region of LotC (**Fig. 4D**). At the core of the interface, an extended loop region of the LotA_N_ insertion domain slots L128 into a shallow pocket of Ub^dist^ formed between I36, L71, and T7 (**Fig. 4E**). Additional ionic interactions are observed around the periphery, including hydrogen bonds to both R72 and R74 of Ub^dist^ as the C-terminus enters the LotA_N_ active site (**Fig. 4E**). This interface appears to play a minor role in the binding of full K6 diUb, as an L128A mutation had minimal effect, and only a severe L128R mutation that introduces a steric clash with the Ub^dist^ I36 patch significantly disrupted LotA_N_ cleavage of the fluorescent K6 diUb substrate (**Fig. 4F**).

The LotA_N_ S1’ site orients K6 of Ub^prox^ into the active site through interactions with the Ub I36 and F4 hydrophobic patches (**Fig. 4A, C**). The human DUB USP30 also recognizes the F4 patch of a K6-linked Ub^prox^, but in a very unique manner involving a ∼100-degree twist (**Fig. 4G, S4D**). In the LotA_N_ structure, Ub^prox^ is sandwiched between the core OTU and insertion domains and makes a single hydrogen bond to R74 of Ub^dist^ through the carbonyl oxygen of T9 (**Fig. 4H**). The F4 side chain of Ub^prox^ makes Van der Waals interactions to the peptide backbone of D10 within the LotA_N_ OTU core domain (**Fig. 4H**). At the Ub^prox^ I36 patch, F120 and L141 from the LotA_N_ insertion domain interact with I36, while buried within the interface lies a salt bridge between LotA_N_ R145 and Ub^prox^ E34. Mutation of these residues in LotA_N_ had a much more severe impact on K6 diUb cleavage than mutations in the S1 site (**Fig. 4F**), as either an F120A mutation to disrupt the hydrophobic interface or an R145L mutation to eliminate the salt bridge caused substantial reduction in DUB activity (**Fig. 4I**). Reciprocal mutations specifically introduced into Ub^prox^ of K6 diUb (see Materials and Methods), showed the strongest defect in LotA_N_ cleavage with mutation of F4, a modest effect with I36 mutation, and no effect upon mutation of the unutilized I44 site (**Fig. 4J**).

The LotA_N_ insertion domain makes key contributions to both the S1 and S1’ Ub-binding sites required for DUB activity. With exception of several loops and extended regions, the core structures of the Lot-class insertion domains follow a related underlying 4-helix topology (**Fig. 4K, S4E**). By structural comparison using Dali (Holm, 2020), the insertion domain of LotA_N_ is the most similar to LotC over other Lot-class DUBs (**Fig. S4F**). Based on the Ub- and diUb-bound structures of LotC and LotA_N_, respectively, it is the unique features of the α1-2 region that dictate interactions to Ub (Liu *et al*, 2020) (**Fig. S4F-H**). Modeling and mutagenesis studies implicate the α1-2 regions of LotB and LotA_M_ in mediating important Ub interactions as well (Shin *et al*, 2020; Takekawa *et al*, 2022). Interestingly, a Dali search of the LotA_N_ insertion domain also detected similarity to the VR-1 region of OtDUB, an effector DUB from *Orientia tsutsugamushi* that belongs to the CE clan of cysteine proteases (Berk *et al*, 2020) (**Fig. S4I**). The OtDUB VR-1 region follows the 4-helix structural topology conserved among Lot-class DUBs, and encodes two Ub-binding sites: one at the edge of Helix 3 that forms part of the S1 site to bind Ub^dist^ (**Fig. S4J**), and a second located in the α1-2 region that binds an additional Ub with unusually high affinity (**Fig. S4K**). Unlike the high affinity site in the α1-2 region (**Fig. S4K**), residues at the edge of Helix 3 in OtDUB VR-1 are conserved in the LotA_N_ structure (**Fig. S4J**). Additional binding of Ub at this site in LotA_N_, however, would cause steric clash with the Ub^dist^ observed in our structure (**Fig. S4J**). Because of its important roles in Lot-class DUBs as well as OtDUB, we propose to assign the name <u>A</u>daptive <u>U</u>biquitin-<u>B</u>inding <u>D</u>omain, or “A-UBD” for short, to this common 4-helix substructure, owing to adaptations of the unique α1-2 regions that impart distinctive Ub interactions into a common topological scaffold (**Fig. 4K, S4E**).

### Conformational changes leading to LotA_N_ activation

Comparison of the apo and K6 diUb-bound LotA_N_ structures revealed several interesting changes. First, the complex structure exhibited a 45-degree swing of the A-UBD with respect to the core OTU domain, resulting in a more closed conformation (**Fig. 5A**). This 45-degree swing moves key residues such as L128, F120, and R145 up, forming the S1 and S1’ Ub-binding sites that sandwich K6 diUb against the active site of the core OTU domain (**Fig. 5B, 4E, 4H**). Movement within the hinge region is required for LotA_N_ activity, as a stabilizing A193P mutation within the hinge reduces cleavage of the fluorescent K6 diUb substrate (**Fig. 5C**). Analogous hinge regions can be observed between the core OTU and A-UBDs of other Lot-class DUBs (**Fig. S5**). Comparison of the apo and Ub-bound LotC structures shows a 6-degree swing, in this case toward a more open conformation in the Ub-bound complex (**Fig. S5**). Thus, it is possible that other Lot-class DUBs may also require mobility of the A-UBD for DUB activity. Another dramatic difference between the apo and diUb-bound LotA_N_ structures is in the active site. As highlighted above, the apo LotA_N_ structure portrays a misaligned catalytic triad (**Fig. 3B**), while the active site in the diUb-bound structure has become properly aligned for catalysis (**Fig. 4B**). Coincident with this rearrangement, the so-called “Cys loop” preceding the catalytic cysteine is shifted by over 4Å, causing a rearrangement of the cysteine’s position toward the remaining triad members. Opposing the LotA_N_ Cys loop is F4 of Ub^prox^, which appears to select for this active conformation by creating steric conflict with the conformation observed in the apo structure (**Fig. 5D**). This contact is critical for LotA_N_ activation, as K6 diUb incorporating an F4A mutation into Ub^prox^ cannot be cleaved (**Fig. 4J**). The so-called “His loop” preceding the general base histidine is also shifted between the apo and bound LotA_N_ structures, resulting in a >3Å shift of the histidine alpha carbon (**Fig. 5D**). Opposing this loop is H68 of Ub^prox^, which makes a hydrogen bond to the backbone carbonyl of E235 in the complex structure (**Fig. 5D**). This interaction is also important for LotA_N_ activation, as K6 diUb incorporating an H68A mutation into Ub^prox^ is cleaved less efficiently than wild-type (**Fig. 5E**). Interestingly, the effect of Ub^prox^ H68 on the LotA_N_ His loop relies on both hydrogen bonding and steric effects, as K6 diUb substrates incorporating either a Ub^prox^ H68Q or H68F mutation are both cleaved less efficiently than wild-type (**Fig. 5E**). Thus, Ub binding at the S1’ site in the correct, K6-specific orientation activates the LotA_N_ catalytic center through substrate-assisted catalysis, by which F4 and H68 of Ub^prox^ assist in the proper orientation of the Cys and His loops.

**Figure 5:**
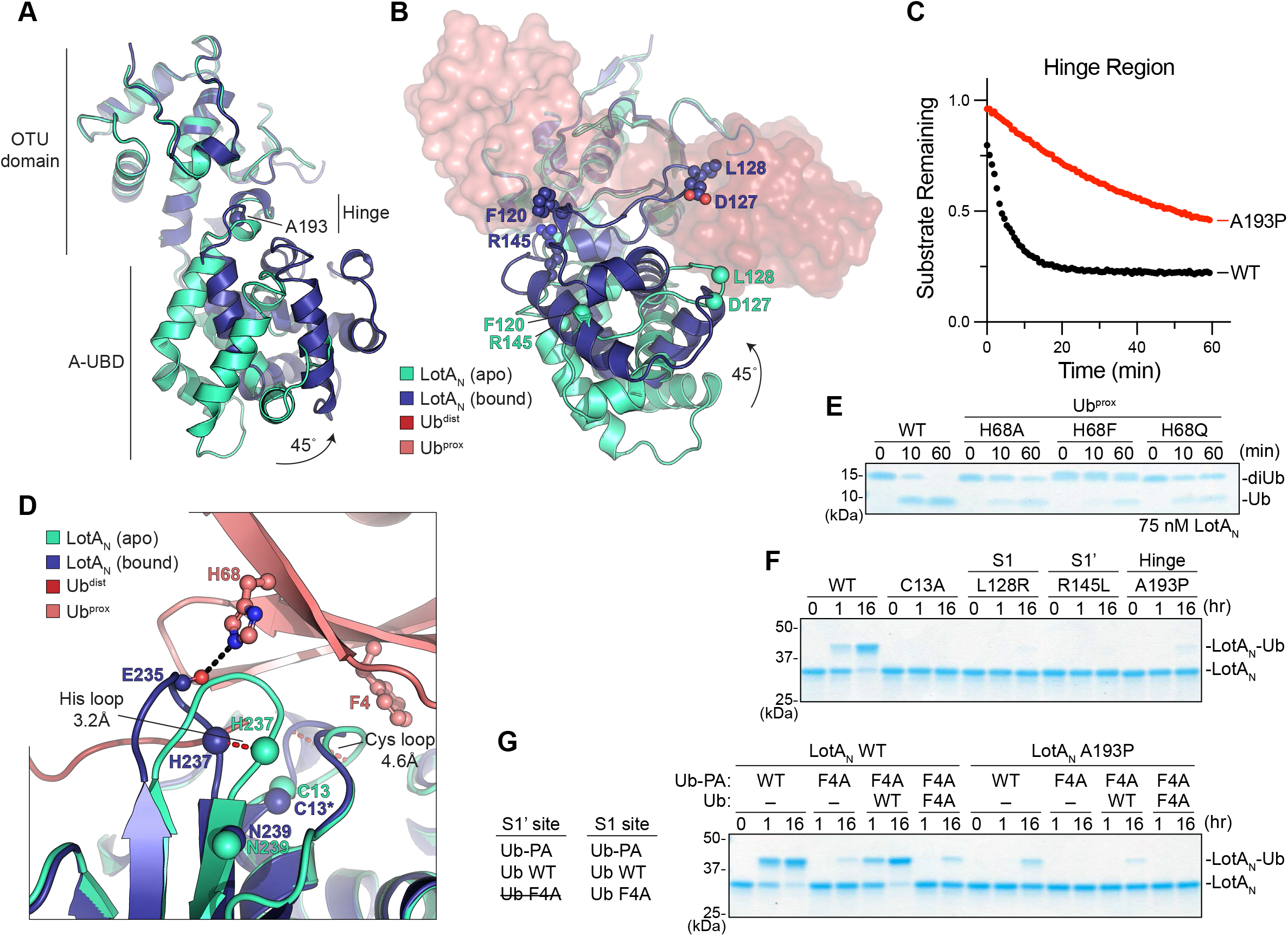
Conformational changes of LotA_N_ during catalysis. A. Structural overlay of apo (cyan) and K6 diUb-bound (blue) LotA_N_, aligned on their core OTU domains. A site for structure-guided mutation within the hinge region is shown. B. As in (**A**), following ∼90-degree rotation to visualize formation of the LotA_N_ S1 and S1’ Ub-binding sites upon rotation of the A-UBD. C. Cleavage of fluorescent K6 diUb by the indicated LotA_N_ hinge variant at 10 nM concentration monitored by fluorescence polarization. These data were collected in parallel with those presented in Fig. 4F and I, and the WT dataset is shown again for reference. D. Structural overlay of the active sites from apo (cyan) and K6 diUb-bound (blue) LotA_N_, aligned on their core OTU domains. Conformational changes of the Cys and His loops that occur upon Ub binding (shades of red) are shown by red dashed lines. Distances of the Cys and His loop movements are measured by the Cα positions of LotA_N_ H237 and G11, respectively. The Cα of LotA_N_ catalytic triad residues are shown as spheres. Ub^prox^ residues opposing the rearranged LotA_N_ loops are shown in ball-and-stick representation, with hydrogen bonds indicated as black dashed lines. E. Gel-based LotA_N_ cleavage assay of K6 diUb variants encoding the indicated Ub^prox^ mutations. Reactions were quenched at the indicated times and visualized by SDS PAGE with Coomassie staining. F. Ub-PA reactivity assay for LotA_N_ wild-type alongside the indicated catalytic, S1 site, S1’ site, and hinge mutations. Reactions were quenched at the indicated times and visualized by SDS PAGE with Coomassie staining. G. Ub-PA reactivity assay combining the indicated LotA_N_, Ub-PA, and Ub variants. The ability of each Ub variant to competently bind the LotA_N_ S1 or S1’ site is listed. *Left*, the importance of the LotA_N_ S1’ site for Ub-PA reactivity was tested by combining F4A Ub-PA, (deficient in binding the S1’ site) with wild-type or F4A Ub as a rescue. *Right*, the importance of LotA_N_ hinge flexibility was tested by performing a matched set of experiments with the A193P hinge mutant. See also Figure S5.

A mechanism of substrate-assisted catalysis is consistent with the observed lack of activity against the monomeric Ub-KG(TAMRA) substrate (**Fig. 1C**), wherein Ub would presumably bind at the S1 site but the attached KG(TAMRA) moiety occupying the S1’ site could not serve to activate the catalytic triad. LotA_N_ can, however, react with the monomeric Ub-PA activity-based probe (**Fig. 1B**). In order to reconcile how Ub-PA binding and reactivity at the S1 site could occur within a mechanism of substrate-assisted catalysis, we tested for Ub-PA reactivity across a series of structure-guided LotA_N_ mutants. As expected, the Ub-PA reactivity observed in wild-type LotA_N_ is abolished or severely reduced in the backgrounds of the catalytic C13A or S1-site L128R mutations, respectively (**Fig. 5F**). Unexpectedly, reactivity was also lost in the background of the LotA_N_ R145L S1’ mutant as well as the A193P hinge mutant (**Fig. 5F**). One hypothesis that could explain this effect is that Ub-PA is able to bind at the S1’ site and activate reactivity for a second Ub-PA molecule bound at the S1 site. This scenario would also explain the lack of activity against Ub-KG(TAMRA), as the bulky KG(TAMRA) moiety would occlude binding of a second Ub molecule into the S1’ site. To test this model, we constructed a Ub-PA probe that carries the F4A mutation, such that it can no longer play a role in activating LotA_N_ at the S1’ site (see **Fig. 4J and 5D**). Compared to incubation with wild-type Ub-PA, the reactivity of LotA_N_ toward the F4A Ub-PA mutant is severely reduced (**Fig. 5G**). Addition of wild-type Ub (lacking the PA warhead) is able to rescue the reactivity of F4A Ub-PA by serving as a dedicated “activator” at the S1’ site (**Fig. 5G**). The analogous rescue experiment with F4A Ub, however, fails to activate LotA_N_ for reactivity with the F4A Ub-PA mutant (**Fig. 5G**). Thus, binding at the S1’ site and fulfillment of substrate-assisted catalysis is a prerequisite for Ub-PA reactivity at the S1 site. Upstream of Ub binding at the S1’ site is flexibility within the LotA_N_ hinge that facilitates a closed conformation competent of Ub binding, as incorporation of the A193P hinge mutant overrides all scenarios of Ub-PA reactivity (**Fig. 5G**).

### LotA_N_ activity is required for restriction of K6 polyUb at the *Legionella*-containing vacuole

LotA, LotB, and LotC have all been shown to edit Ub modifications deposited onto the LCV (Kubori *et al*, 2018; Ma *et al*, 2020; Liu *et al*, 2020). LotA has been shown to play a role in regulating K6, K48, and K63 polyUb at the LCV surface following localization via a PI(3)P-binding domain at its C-terminus (Kubori *et al*, 2018) (**Fig. 1A**). Whether these signal types are independently regulated by LotA’s two OTU domains, however, had not been tested. To test whether the ability of LotA_N_ to restrict K6 polyUb *in vitro* holds true during infection, we constructed *L. pneumophila* strains with genomic *lotA* mutations at either OTU active site (C13S, C303S) for comparison to the wild-type Lp01 and Δ*lotA* strains, and performed infections in HeLa-FcγRII cells. To circumvent the absence of a good detection reagent for K6 polyUb, we transfected the HeLa-FcγRII cells with HA-Ub-K6 (in which all lysines are mutated to arginine except for K6) prior to infection as a reporter for K6 polyUb (see Materials and Methods). Compared to the wild-type strain, deletion of *lotA* results in a significant increase in the number of K6-positive LCVs (**Fig. 6A-B**). Further comparison to the inactivating mutations in each OTU domain reveals that the C303S mutation in LotA_M_ retains the ability to restrict K6 polyUb at wild-type levels, while the LotA_N_ C13S mutant behaves similarly to total *lotA* deletion (**Fig. 6A-B**). These findings are consistent with the differing specificities we observe for LotA_N_ and LotA_M_ *in vitro* (**Fig. 1D-G**), and suggest a delineation for their roles during infection.

**Figure 6:**
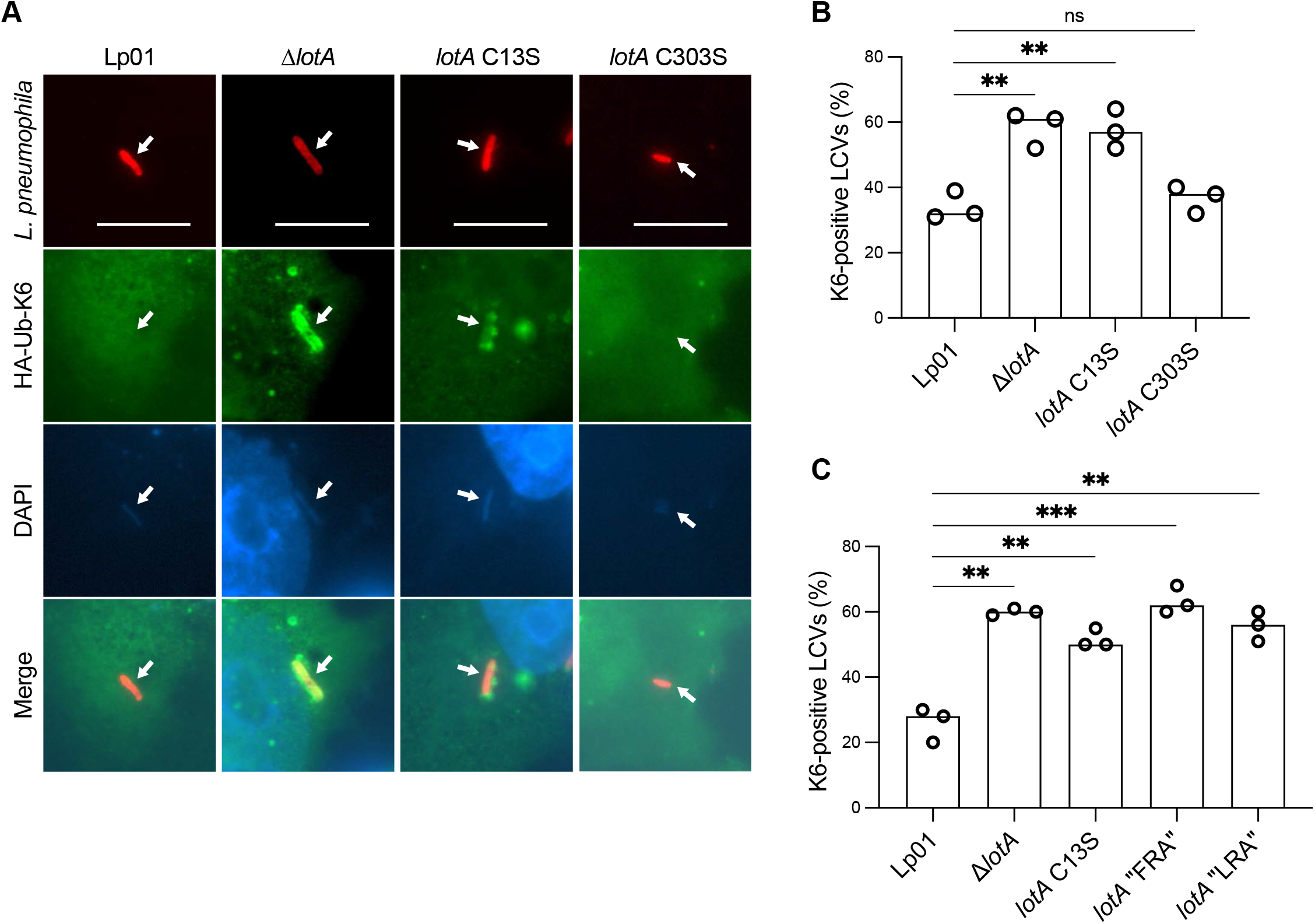
LotA_N_ restriction of K6 polyUb during *L. pneumophila* infection. A. Representative images of HeLa FcγRII cells infected with the indicated *L. pneumophila* strains at an MOI of 2 for 4 h. The C13S mutation renders LotA_N_ inactive, while the C303S mutation renders LotA_M_ inactive. Fixed cells were stained for *L. pneumophila* (red), HA-Ub-K6 (green), and DNA (blue). Arrows indicate the position of a bacterium in each channel. Scale bars correspond to 10 µm. B. Quantitation of K6-positive bacteria shown in (**A**). Infections were performed in triplicate and each value represents scoring from 100 LCVs. Significance was determined using a Welch’s t-test. C. Quantitation of K6-positive bacteria from HeLa FcγRII cells infected with the indicated *L. pneumophila* strains at an MOI of 2 for 4 h. The “FRA” triple mutant combines the LotA_N_ S1’-site F120A, S1’-site R145L, and hinge A193P mutations. The “LRA” triple mutant combines the LotA_N_ S1-site L128R, S1’-site R145L, and hinge A193P mutations. Infections were performed in triplicate and each value represents scoring from 100 LCVs. Significance was determined using a Welch’s t-test. See also Figure S6.

As a further test of our structure-guided mechanistic studies, we sought to examine the effects of mutations outside of the catalytic cysteine on the ability of LotA_N_ to restrict K6 polyUb at the LCV. We designed two separate triple-mutant variants of LotA_N_: “FRA” combines the S1’-site mutations F120A and R145L with the hinge mutation A193P, while “LRA” combines the S1-site mutation L128R, the S1’ mutation R145L, and the hinge mutation A193P. *In vitro*, neither of the triple-mutant variants show any detectable cleavage of the fluorescent K6 diUb substrate, even at high concentration (**Fig. S6A**). *L. pneumophila* strains carrying either of the *lotA* triple-mutant variants failed to restrict K6 polyUb on the LCV to a similar extent as the *lotA* C13S or complete *lotA* deletion strains (**Fig. 6C, S6B**), demonstrating that the requirements for LotA_N_ DUB activity *in vitro* are also present during infection.

## DISCUSSION

By separating the two OTU domains of LotA, we have revealed interesting distinctions in their DUB activities and specificities. While the second OTU domain, LotA_M_, exhibits activity toward longer K48, K6, and K63 polyUb, the first OTU domain, LotA_N_, is extremely specific and highly active toward K6 polyUb. The identification of K6-specific enzymes in humans has proven difficult. Many ligases and DUBs that regulate K6 linkages also demonstrate considerable activity toward other types of polyUb (Ordureau *et al*, 2014; Michel *et al*, 2017; Gersch *et al*, 2017; Sato *et al*, 2017; Mevissen *et al*, 2013). Perhaps coincidentally, the most K6-specific E3 ligase identified to date, NleL, also originates from bacteria (Lin *et al*, 2011). Just as NleL has become an invaluable tool for studying K6 polyUb (Hospenthal *et al*, 2013), LotA_N_ will be equally useful, particularly in applications such as UbiCRest analyses. Prior use of OTUD3 as a K6-targeted DUB has required careful consideration of enzyme concentration and secondary activities (Hospenthal *et al*, 2015). We show that LotA_N_ maintains K6 specificity even at high concentration against long, complex polyUb chains, and will thus be a convenient tool to be used along with other polyUb-specific DUBs.

The exquisite specificity of LotA_N_ toward K6 polyUb arises from a unique blend of regulatory mechanisms (**Fig. 7**). In our apo structure of LotA_N_, the open conformation of the OTU domain and A-UBD is incompatible with polyUb binding and, in addition, the active site is not in the proper alignment for catalysis. Consistent with inherent flexibility in these regions, the LotA_N_ Cys loop, His loop, and hinge region all exhibit above average B-factors. Before polyUb binding can occur, flexibility is required in the LotA_N_ hinge region in order for the A-UBD to swing into position. This *in situ* formation of the S1/S1’ Ub-binding sites resembles the conformational changes that occur as the human OTU DUB Cezanne binds to K11 diUb (Mevissen *et al*, 2016). Once LotA_N_ occupies a more closed OTU:A-UBD arrangement, the unique topology of K6 diUb allows it to bind and properly align the catalytic triad. Only a K6-linked Ub^prox^ would present the proper configuration of F4 and H68 to select for the catalytically-competent alignment of the Cys and His loops, respectively. The human DUB OTULIN uses a similar mechanism of substrate-assisted catalysis, by which the unique position of a residue from an M1-linked Ub^prox^ is required for alignment of the catalytic triad (Keusekotten *et al*, 2013). Thus, LotA_N_ has been evolved to encode multiple layers of regulation that act in concert to maintain K6 polyUb specificity. Unlike LotB and LotC, LotA_M_ also appears to require some form of substrate-assisted catalysis for active site rearrangement, in addition to its requirement for longer polyUb chains.

**Figure 7:**
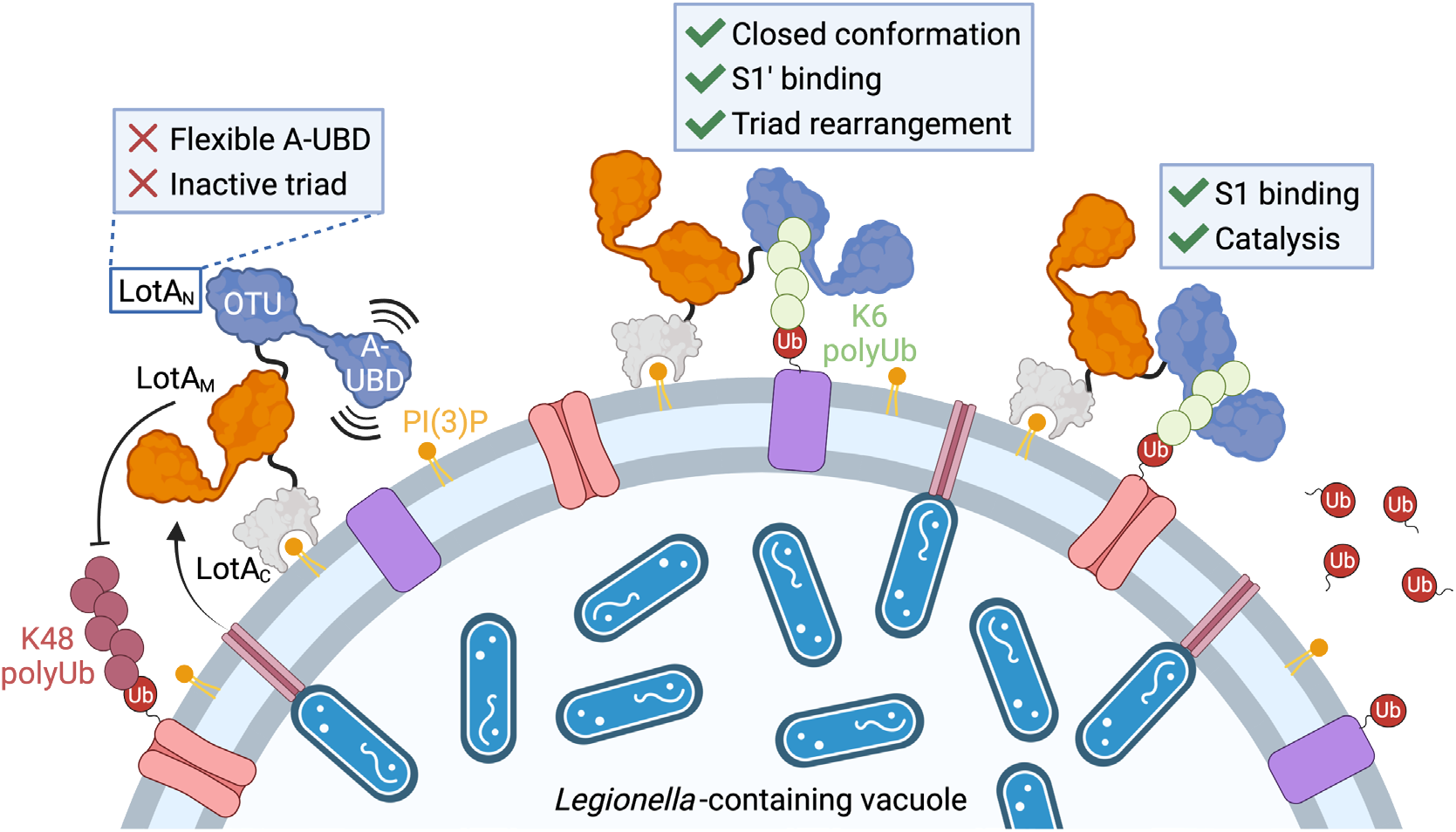
Model for LotA deubiquitinase activity. Secreted LotA localizes back to the cytosolic face of the LCV via the PI(3)P-binding LotA_C_ domain. At the LCV, DUB activity of LotA_M_ acts to restrict long K48 and K63 polyUb. LotA_N_ is kept inactive by a flexible A-UBD and inactive arrangement of the catalytic triad. Formation of a closed OTU:A-UBD conformation allows formation of an S1’ Ub-binding site. Binding of a K6-linked Ub into the S1’ site orients the LotA_N_ catalytic triad, and allows for hydrolysis of K6 polyUb. Illustration created with BioRender.com.

Helical sub-domains of approximately 100 amino acids in length inserted into VR-1 of the OTU domain are one of the defining features of Lot-class DUBs. While the basic ∼4-helix fold is similar across the Lot-class structures reported thus far, adaptations to the α1-2 region add versatility to the Ub interface, hence our proposed name <u>A</u>daptive <u>U</u>biquitin-<u>B</u>inding <u>D</u>omain, or “A-UBD” (**Fig. 4K, S4E**). The α1-2 regions of the A-UBDs encoded by LotA_N_, LotA_M_, LotB, LotC, as well as OtDUB have all be shown to play important roles in mediating Ub interactions (Takekawa *et al*, 2022; Shin *et al*, 2020; Liu *et al*, 2020; Berk *et al*, 2020). It is remarkable that this helical substructure, now observed in at least 5 DUB domains across two bacterial species, has adopted such unique modes of Ub interaction. To what extent the A-UBD is incorporated into other regulators of Ub signaling across bacteria, viruses, and eukaryotes will be an interesting area of future research.

During infection, LotA has been shown to localize to the LCV via its C-terminal PI(3)P-binding domain, where it acts to restrict K6, K48, and K63 polyUb signals (Kubori *et al*, 2018). Through separation of the two OTU domains, we show that not only do LotA_N_ and LotA_M_ display distinct specificities toward polyUb substrates, but they also regulate distinct polyUb signals on the LCV surface (**Fig. 7**). LotA_N_ activity is required for LotA-dependent restriction of K6 signals. The close spacing of the two LotA domains (∼20 amino acids), however, raises the interesting possibility that they work cooperatively on a large or particularly complex polyUb signal. The evolution of such a K6-specific DUB by *L. pneumophila* also raises interesting questions as to what cellular process LotA_N_ acts to restrict. K6 polyUb signals have previously been tied to mitophagy, i.e., the autophagic destruction of damaged mitochondria, where they are thought to primarily be regulated by the E3 ligase Parkin and DUB USP30 (Swatek & Komander, 2016). Interestingly, Parkin has also been implicated in the Ub-mediated autophagic destruction of intracellular bacteria such as *Mycobacterium tuberculosis* (Manzanillo *et al*, 2013). Among its many secreted effector proteins, *L. pneumophila* has evolved several activities that act to restrict host autophagic defenses (Thomas *et al*, 2020). Whether and how LotA_N_ might fit into this network, and what additional facets of K6 polyUb biology might be uncovered as a result, will be an exciting area of future work.

## MATERIALS AND METHODS

### Cloning and mutagenesis

The *lotA* gene was cloned from *Legionella pneumophila* Philadelphia-1 genomic DNA. Initial construct design was performed using Phyre2 (Kelley *et al*, 2015) and the LotA_M_ crystal structure (Takekawa *et al*, 2022), yielding construct boundaries of 1-300 and 287-543 for LotA_N_ and LotA_M_, respectively. To support crystallization, additional C-terminal truncations of LotA_N_ led to the 1-294 construct used for determination of the apo LotA_N_ structure. Based on the apo structure, an additional LotA_N_ construct encoding amino acids 1-276 was used for determination of the K6 diUb-bound structure. All LotA constructs were cloned into the pOPINB vector, which encodes an N-terminal His tag followed by a 3C protease cleavage site. Cloning and mutagenesis were performed using Phusion DNA Polymerase (New England BioLabs) and TOP10 *Escherichia coli* (MilliporeSigma).

### Protein expression and purification

All pOPINB-LotA constructs were expressed and purified similarly. Transformed Rosetta (DE3) *Escherichia coli* were grown in Luria broth containing 35 mg/mL chloramphenicol and 50 mg/mL kanamycin at 37°C until an optical density (600 nm) of 0.6-0.8. Cultures were cooled to 18°C and induced with 200 mM IPTG for an additional 20 hours. Cells were harvested by centrifugation and resuspended in 25 mM Tris, 200 mM NaCl, 2 mM β-mercaptoethanol, pH 8.0 (Buffer A). Following a freeze-thaw cycle, cells were incubated on ice with lysozyme, DNase, and SigmaFAST protease inhibitor cocktail (MilliporeSigma) prior to lysis by sonication. Clarified lysates were applied to HisPur cobalt affinity resin (ThermoFisher) and washed with Buffer A prior to elution with Buffer A containing 300 mM imidazole. Eluted proteins were cleaved with 3C protease overnight during dialysis back to Buffer A at 4°C. Cleaved LotA proteins were concentrated using 10K MWCO Amicon centrifugal filters (MilliporeSigma) and applied to a HiLoad Superdex 75 pg size exclusion chromatography column (Cytiva) equilibrated in 25 mM Tris, 200 mM NaCl, 5 mM DTT, pH 8.0. Fractions were evaluated for purity by SDS PAGE, collected, concentrated, and quantified by absorbance (280 nm) prior to flash freezing and storage at −80°C.

Untagged Ub constructs were expressed from the pET-17b vector. Transformed Rosetta (DE3) *Escherichia coli* were grown by auto-induction in a modified ZYM-5052 media (Studier, 2005) containing 35 mg/mL chloramphenicol and 100 mg/mL ampicillin at 37°C for 24-48 h. Cells were harvested by centrifugation, resuspended, and lysed as above for LotA. Clarified lysates were acidified by dropwise addition of 70% perchloric acid to a final concentration of 0.5%. The mixture was allowed to stir on ice for 1-2 h prior to centrifugation. The supernatant was dialyzed against 50 mM sodium acetate, pH 4.5 overnight at 4°C. The protein was applied to a HiPrep SP FF 16/10 cation exchange column (Cytiva), washed with additional 50 mM sodium acetate, pH 4.5, and eluted over a linear gradient to a matched buffer containing 500 mM NaCl. Fractions were evaluated for purity by SDS PAGE, pooled, and dialyzed against 25 mM Tris, 200 mM NaCl, pH 8.0 overnight at 4°C. If necessary, further purification was performed using a HiLoad Superdex 75 pg size exclusion chromatography column (Cytiva) equilibrated in 25 mM Tris, 200 mM NaCl, pH 8.0. Purified Ub was concentrated using 3K MWCO Amicon centrifugal filters (MilliporeSigma), quantified by absorbance (280 nm), and flash frozen for storage at either - 20°C or −80°C.

The Ub-PA activity-based probes were prepared using intein chemistry as described previously (Wilkinson *et al*, 2005). Rosetta (DE3) *Escherichia coli* transformed with pTXB1-Ub(1-75) were grown in Luria broth containing 35 mg/mL chloramphenicol and 100 mg/mL ampicillin at 37°C until an optical density (600 nm) of 0.6-0.8. Cultures were cooled to 18°C and induced with 500 mM IPTG for an additional 20 hours. Cells were harvested by centrifugation and resuspended in 20 mM HEPES, 50 mM sodium acetate, 75 mM NaCl, pH 6.5 (Buffer B). After a freeze-thaw cycle, cells were incubated on ice with DNase, SigmaFAST protease inhibitor cocktail (MilliporeSigma), and PMSF prior to lysis by sonication. The clarified lysate was applied to chitin resin (New England BioLabs) and allowed to bind at 37°C for 2 h. The resin was washed with Buffer B, followed by a high salt wash with Buffer B containing 500 mM NaCl, followed by a final wash with Buffer B. The Ub-MesNa was allowed to form and elute from the chitin resin for 42 h at room temperature, following addition of Buffer B containing 100 mM MesNa. The eluted protein was concentrated using 3K MWCO Amicon centrifugal filters (MilliporeSigma) and applied to a HiLoad Superdex 75 pg size exclusion chromatography column (Cytiva) equilibrated in Buffer B. Fractions containing Ub-MesNa were evaluated for purity by SDS PAGE and pooled. To convert Ub-MesNa to Ub-PA, propargylamine hydrochloride was added in 1000-fold excess and, following adjustment of the pH to 8.5, incubated at room temperature for 3 h. The reaction was dialyzed against 25 mM Tris, 200 mM NaCl, pH 8.0 overnight at 4°C. The final product was concentrated using 3K MWCO Amicon centrifugal filters (MilliporeSigma), quantified by absorbance (280 nm), and stored on ice during immediate use.

### Assembly of polyUb chains

All diUb and tetraUb chains were assembled and purified according to published methods (Michel *et al*, 2018), with the exception of K27 diUb and K11 tetraUb, which were purchased from UbiQ and R&D Systems, respectively. K6 diUb for fluorescent labeling or for experiments that incorporate Ub^prox^ mutations were assembled using a “capped” strategy that allows directed formation of K6-linked diUb from dedicated donor and acceptor Ub moieties that are incorporated as Ub^dist^ and Ub^prox^, respectively. The dedicated donor Ub construct incorporated K6R and K48R mutations, but retained an intact C-terminus. The dedicated acceptor Ub construct incorporated a K48R mutation and lacked the C-terminal LRGG motif. Effects of Ub^prox^ mutation were assessed by incorporating additional mutations into the dedicated acceptor Ub construct. Alternatively, a Ub^prox^ for fluorophore labeling was incorporated with an acceptor Ub construct that coupled a deletion of G76 with a G75C mutation. Equimolar amounts of donor and acceptor Ub variants were combined with 100 nM UBE1, 600 nM UBE2L3, 10 µM NleL, and 10 mM ATP in buffer containing 40 mM Tris, 10 mM MgCl_2_, 0.5 mM DTT, pH 8.5. Reactions were incubated at 37°C for 16 h and subsequently quenched by addition of 10 mM DTT and 50-fold dilution into 50 mM sodium acetate, pH 4.5. The reactions were applied to a HiPrep SP FF 16/10 cation exchange column (Cytiva), washed with additional 50 mM sodium acetate, pH 4.5, and eluted over a linear gradient to a matched buffer containing 500 mM NaCl. Fractions were evaluated for purity by SDS PAGE, pooled, and dialyzed against 25 mM Tris, 200 mM NaCl, pH 8.0 overnight at 4°C. The diUb product was concentrated using 3K MWCO Amicon centrifugal filters (MilliporeSigma), quantified by absorbance (280 nm), flash frozen, and stored at either −20°C or −80°C.

Fluorescent K6 diUb was prepared using the “capped” strategy described above, which allows specific labeling of a cysteine incorporated at position 75 of Ub^prox^. Purified K6 diUb was diluted to 50 µM in 25 mM sodium phosphate, 150 mM NaCl, 10 mM DTT, pH 7.4 and incubated at room temperature for 30 min. The diUb was subsequently buffer exchanged into 25 mM sodium phosphate, 150 mM NaCl, pH 7.4 over a PD-10 desalting column (Cytiva). ATTO488-maleimide (ATTO-TEC) was added in 5-fold molar excess to 1 mL of the desalted K6 diUb, and incubated at room temperature in the dark for 18 h. The labeled K6 diUb was again desalted to remove excess fluorophore, quantified by absorbance (488 nm), and flash frozen for storage at − 80°C.

### Gel-based DUB assays

Gel-based DUB assays were performed according to published methods (Pruneda & Komander, 2019). DUBs were prepared at 2X concentration in 25 mM Tris, 150 mM NaCl, 10 mM DTT and incubated at room temperature for 10 min. Equal volumes of 2X DUB and either 10 µM diUb or 5 µM tetraUb were mixed and incubated at 37°C. Samples were quenched in sample buffer and analyzed by SDS PAGE.

### Ub-PA reactivity assays

Ub-PA reactivity assays were performed according to published methods (Pruneda & Komander, 2019). DUBs were diluted to 10 µM in 25 mM Tris, 150 mM NaCl, 10 mM DTT, pH 7.4 and incubated at room temperature for 10 min. Equal volumes of 2X DUB and 100 µM Ub-PA were mixed and allowed to react at room temperature for up to 18 h, prior to being quenched in sample buffer and analyzed by SDS PAGE. For rescue experiments with the F4A Ub-PA variant, 25 µM wild-type or F4A Ub was added to the above reaction.

### Fluorescence-based DUB assays

Ub- and Ubl-KG(TAMRA) substrates were prepared as previously described (Geurink *et al*, 2012; Basters *et al*, 2014). Cleavage of the Ub- and Ubl-KG(TAMRA) substrates was monitored by fluorescence polarization according to published methods (Pruneda & Komander, 2019). Enzymes and substrates were prepared at 2X concentration in 25 mM Tris, 100 mM NaCl, 5 mM β-mercaptoethanol, 0.1 mg/mL BSA, pH 7.4 and mixed 1:1 to initiate the reaction. All substrates were held at 50 nM concentration, while protease concentrations were adjusted according to their specific activities. Fluorescence polarization was recorded using a BMG LabTech ClarioStar plate reader with an excitation wavelength of 540 nm, an LP 566 nm dichroic mirror, and an emission wavelength of 590 nm. Reactions were performed in Greiner 384-well small-volume HiBase microplates with 20 μL reaction volumes. Three technical replicates were performed for each reaction, with additional wells containing KG(TAMRA) and Ub/Ubl-KG(TAMRA) as positive and negative controls, respectively. Fluorescence polarization was recorded once per minute over a 1-2 h time period. Data collected from the positive and negative control wells were used to convert fluorescence polarization measurements to fraction of substrate remaining.

DUB assays using the ATTO488-labeled K6 diUb substrate were performed as above, with minor changes. Fluorescence polarization was monitored using an excitation wavelength of 482 nm, an LP 504 nm dichroic mirror, and an emission wavelength of 530 nm. Assay concentrations were 10 nM LotA_N_ and 50 nM ATTO488-labeled K6 diUb, except for the LotA_N_ triple-mutants, which were also assayed at a higher enzyme concentration of 1 µM. ATTO488-labeled monoUb and diUb were used as positive and negative controls, respectively, and used to convert fluorescence polarization measurements to fraction of substrate remaining. To determine Michaelis-Menten parameters, LotA_N_ concentration was held at 4 nM and initial rates of cleavage were determined from the linear range of data produced over the first 15-25 minutes of the reaction, across a range of substrate concentrations from 100 nM to 50.1 µM. All substrate concentrations maintained a fixed, 100 nM concentration of ATTO488-labeled K6 diUb, while unlabeled K6 diUb (shown to be hydrolyzed equally well) was used for the remainder. Three technical replicates were performed for each reaction, and three separate trials were performed for each K6 diUb concentration. Positive and negative control measurements were used to convert fluorescence polarization data to amount of substrate consumed, which was used to derive initial rates. Prism 9 was used for Michaelis-Menten analysis of the initial rate data.

### UbiCRest analysis

Linkage-specific polyUb chains were assembled according to published methods (Michel *et al*, 2018). NleL and HUWE1 chain assemblies were performed at 37°C for 2 h in 25 mM Tris, 150 mM NaCl, 10 mM MgCl_2_, 0.5 mM DTT, pH 7.4 using 375 nM UBE1, 1.5 µM Lys-less UBE2L3, 5 mM ATP, and either 3.75 µM NleL (aa 170-782) or 15 µM HUWE1 (aa 3993-4373). Prior to DUB treatment, all polyUb assembly reactions were quenched by addition of 50 mM EDTA and 5 mM DTT. DUBs were diluted to 10 µM in 25 mM Tris, 150 mM NaCl, 10 mM DTT and incubated at room temperature for 10 min. Equal volumes of polyUb assembly and DUB were mixed and incubated at 37°C for 2 h, prior to quenching in sample buffer and analysis by SDS PAGE.

### Protein crystallization and structure determination

LotA_N_ (1-294), LotA_N_ (1-276), and K6 diUb were prepared as described above and exchanged into 25 mM Tris, 125 mM NaCl, 5 mM DTT, pH 7.4. LotA_N_ (1-294) was concentrated to 12 mg/mL and crystallized in hanging drop format with 30% PEG 2K MME, 0.2 M KI, 0.1 M MES pH 6.5 at 20°C in a 1 µL drop with 1:1 protein:precipitant ratio. LotA_N_ (1-276) was mixed with an equimolar amount of K6 diUb and concentrated using a 3K MWCO Amicon centrifugal filter (MilliporeSigma) to a total protein concentration of 8 mg/mL (determined by Bradford Assay). The protein complex was crystallized in hanging drop format with 25% PEG 3350, 10% ethylene glycol, 0.2 M KSCN, 0.1 M MES pH 5.5 at 20°C in a 1 µL drop with 1:1 protein:precipitant ratio. Apo LotA_N_ crystals were cryoprotected in mother liquor containing 12.5% glycerol, while complex LotA_N_ crystals were cryoprotected in mother liquor containing 25% glycerol prior to vitrification.

Diffraction data were collected at the Stanford Synchrotron Radiation Lightsource (SSRL), beamline 9-2. Images were integrated using XDS (Kabsch, 2010) and scaled using Aimless (Evans & Murshudov, 2013). The apo LotA_N_ (1-294) structure was determined by SAD using the SHELXC/D/E pipeline in CCP4i2 (Sheldrick, 2010; Potterton *et al*, 2018), based on anomalous signal arising from bound iodide ions that were present in the crystallization condition. Automated model building was performed using ARP/wARP (Langer *et al*, 2008), followed by iterative rounds of manual model building in COOT and refinement in PHENIX (Emsley *et al*, 2010; Adams *et al*, 2010). The complex LotA_N_ (1-276) structure bound to K6 diUb was determined by molecular replacement with Phaser in CCP4i2, using the apo LotA_N_ and Ub (PDB 1UBQ) structures (McCoy *et al*, 2007; Potterton *et al*, 2018; Vijay-Kumar *et al*, 1987). Model building and refinement were performed as above using COOT and PHENIX (Emsley *et al*, 2010; Adams *et al*, 2010). All figures were generated using PyMOL (www.pymol.org).

### Legionella strains and culture

*L. pneumophila* strains used in this study are listed in Table S2. Point mutations in genomic *lotA* were introduced by a gene knock-in method, whereby mutated *lotA* genes were integrated into the Δ*lotA* strain at the endogenous locus (Kubori *et al*, 2018). Plasmids used for the integration were constructed based on pSR47S-Δ*lotA* (Kubori *et al*, 2018), which contains homologous regions upstream and downstream of the endogenous *lotA* locus inserted into the pSR47S gene-replacement vector as follows (Merriam *et al*, 1997). The linearized pSR47S-Δ*lotA* was generated by PCR using primers lpg2248KI-1 and lpg2248KI-2. Fragments of the mutated *lotA* genes were obtained by PCR using primers lpg2248KI-3 and lpg2248KI-4 from pMMB207-3xFLAG-LotA C13S (Kubori *et al*, 2018), 3xFLAG-LotA C303S (Kubori *et al*, 2018), 3xFLAG-LotA C13S/C303S (Kubori *et al*, 2018), pOPINB-LotA F120A/R145L/A193P, and pOPINB-LotA L128R/R145L/A193P. The vector and *lotA* fragments were ligated using Gibson assembly, and the resulting plasmids were transformed into *E. coli* DH5αλpir. The allelic exchange was conducted as described previously (Zuckman *et al*, 1999). The genomic mutations of *lotA* were validated by PCR and subsequent DNA sequencing. The *L. pneumophila* strains were grown at 37°C in liquid N-(2-acetamido)-2-aminoethanesulfonic acid (ACES; MilliporeSigma)-buffered yeast extract (AYE) media or on charcoal-yeast extract (CYE) plates, as described previously (Horwitz, 1983).

### Cell culture

HeLa-FcγRII cells were grown in Minimum Essential Medium α (MEMα; Gibco) supplemented with 10% fetal bovine serum (FBS) (Arasaki *et al*, 2017). To maintain FcγRII expression, 400 µg/mL hygromycin (Nacarai) was supplied in the media.

### Transfection and infection

HeLa-FcγRII cells were seeded on cover slips in 24-well tissue culture plates at 5 × 10^4^ cells per well 24 h before transfection. Transfection was performed using Lipofectamine2000 (Invitrogen) and 500 ng of pRK5-HA-Ub-K6 (Addgene, #22900) for 24 h according to the manufacturer’s recommendation. Because of the lysine-to-arginine mutations and the encoded N-terminal epitope tag, HA-Ub-K6 can only be utilized for a) monoubiquitination, b) as a distal capping Ub on the end of a polyUb chain, or c) incorporation into a K6-linked chain. Owing to its stringent requirement for substrate-assisted catalysis, LotA_N_ would only be able to hydrolyze HA-Ub-K6 that is incorporated into K6-linked chains. For infection, *L. pneumophila* was cultivated for 24 h in AYE media (with starting A_600_ of 0.2) at 37°C with shaking. The cells were infected with the *Legionella* strains opsonized with anti-*Legionella* rat antibody (Scrum, #17457; 1:3,000 dilution) at a multiplicity of infection (MOI) of 2. After adding bacteria to the cells, the plates were centrifuged at 200 × g to precipitate bacteria onto the layer of cells and were immediately warmed in a 37°C water-bath by floating for 5 min and then placed in a CO_2_ incubator at 37°C. At 1 h after infection, the infected cells were washed three times with prewarmed Dulbecco’s Phosphate Buffered Saline (DPBS; MilliporeSigma) to remove the extracellular bacteria and refreshed with prewarmed media, then incubation was resumed at 37°C in a CO_2_ incubator. At 4 h after infection, the cells were washed three times with DPBS and fixed with 4% (w/v) paraformaldehyde for 20 min.

### Immunofluorescent microscopy

The fixed HeLa-FcγRII cells on coverslips were permeabilized by treating with cold methanol for 60 sec and washed three times with DPBS. The samples were blocked with 2% (v/v) goat serum in DPBS and treated with anti-HA rabbit antibody (MBL, #561) with 1:200 dilution for 90 min. After three times washing with DPBS, the samples were treated with anti-*Legionella* rat antibody (Scrum, #17457; 1:3,000 dilution) for 60 min. After three times washing with DPBS, the samples were treated with secondary fluorescent antibodies, Alexa 488 goat anti-rabbit IgG (Invitrogen, #A11034; 1:500 dilution) and Alexa 568 goat anti-rat IgG (Invitrogen, #A11077; 1:500 dilution), for 40 min. After three times washing with DPBS, the coverslips were rinsed with distilled water and mounted on glass slides with ProLong^TM^ Diamond Antifade Mountant with DAPI (Invitrogen, P36962). Images were collected with an inverted microscope (TE2000-U; Nikon) equipped with a digital ORCA-ERA camera (Hamamatsu).

### Statistical analysis

Welch’s t tests were performed using Prism 9 software with data from three independent experiments.

## ACKNOWLEDGEMENTS

We thank R. Klevit (University of Washington), D. Komander (Walter and Eliza Hall Institute of Medical Research), and T. Mund (MRC Laboratory of Molecular Biology) for sharing expression plasmids, and K. Arasaki (Tokyo University of Pharmacy and Life Sciences) for sharing the HeLa-FcγRII cell line. We thank members of our laboratories and the Seattle Ub Research Group for helpful discussions. Use of the Stanford Synchrotron Radiation Lightsource, SLAC National Accelerator Laboratory, is supported by the U.S. Department of Energy, Office of Science, Office of Basic Energy Sciences under Contract No. DE-AC02-76SF00515. The SSRL Structural Molecular Biology Program is supported by the DOE Office of Biological and Environmental Research, and by the National Institutes of Health, National Institute of General Medical Sciences (P30GM133894). The contents of this publication are solely the responsibility of the authors and do not necessarily represent the official views of NIGMS or NIH. This work was supported by the Takeda Science Foundation (TKu), MEXT/JSPS KAKENHI grants (19H03469 to TKu), Oregon Health & Science University (JNP), The Collins Medical Trust (JNP), and the NIGMS (R35GM142486 to JNP).

## AUTHOR CONTRIBUTIONS

Conceptualization, JNP; Investigation, GDW, TKi, TGF, JVN, TKu, and JNP; Resources, PPG; Writing – original draft, GDW and JNP; Writing – review & editing, GDW and JNP; Supervision, TKu, HN, and JNP; Funding acquisition, TKu and JNP.

## DATA AVAILABILITY

Coordinates and structure factors for the apo and K6 diUb-bound LotA_N_ structures have been deposited in the Protein Data Bank under accession codes 7UYG and 7UYH, respectively. All other data are available upon request.

---

**Supplementary Figure 1:**
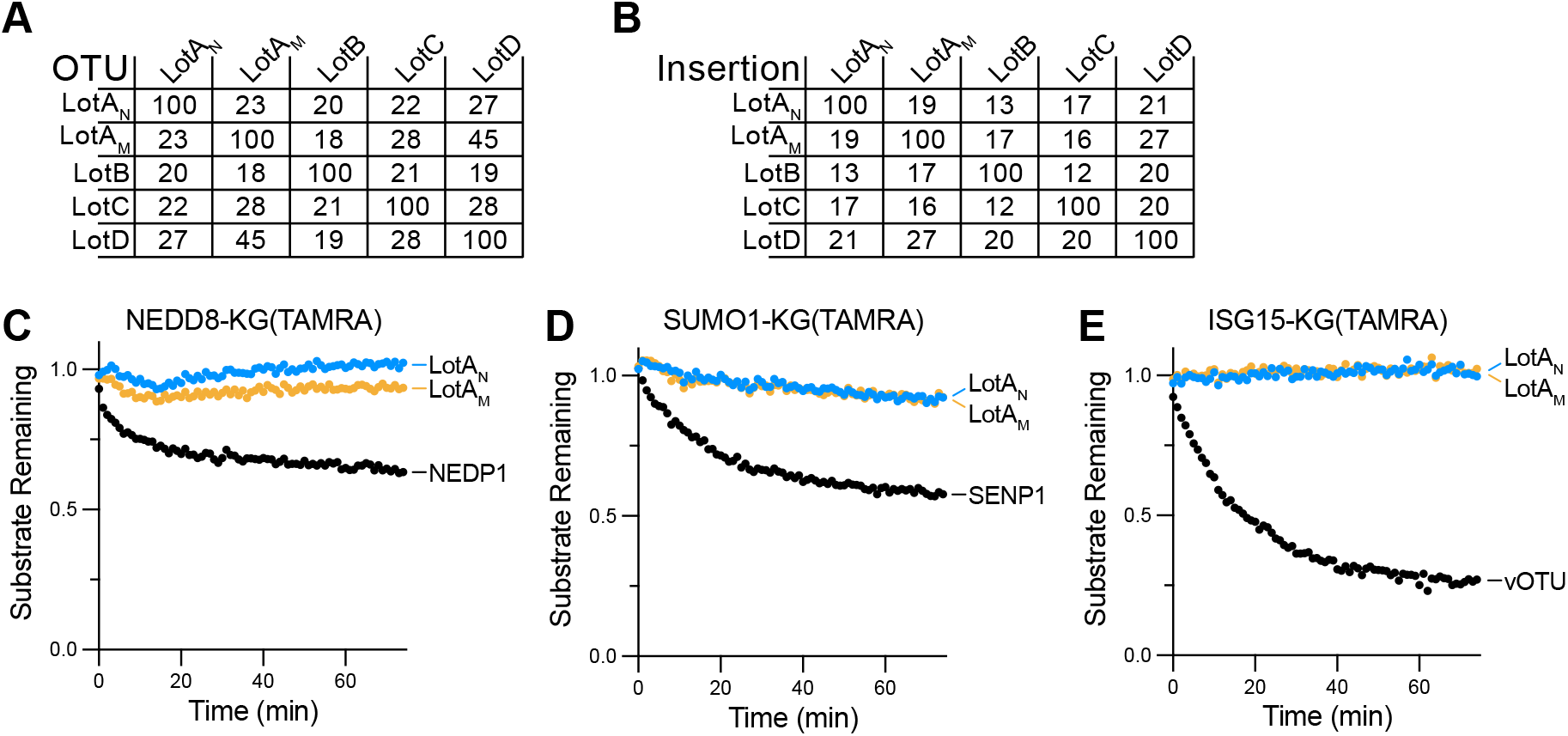
Separation of LotA deubiquitinase activities. A. Pairwise sequence identity matrix calculated from a T-Coffee multiple sequence alignment (Notredame *et al*, 2000) performed on the Lot-class core OTU domain sequences following structure-guided removal of the insertion domains. B. As in (**A**), for the Lot-class insertion domains. C-E. Ub-like-KG(TAMRA) cleavage assays for NEDD8 (**C**), SUMO1 (**D**), and ISG15 (**E**) monitored by fluorescence polarization. Activity was measured using 1 µM LotA_N_, 1 µM LotA_M_. 6 nM NEDP1, 250 pM SENP1, and 400 nM vOTU were used as control enzymes for NEDD8, SUMO1, and ISG15, respectively.

**Supplementary Figure 2:**
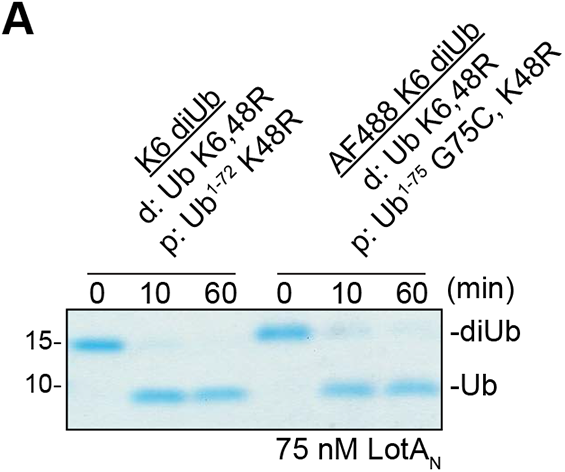
Application of LotA_N_ K6 specificity for UbiCRest analysis. A. Gel-based LotA_N_ cleavage assay comparing the capped and fluorescent capped K6 diUb substrates used in the kinetic analysis. Capped K6 diUb incorporated the annotated mutations into the distal (d) and proximal (p) Ub moieties to streamline production.

**Supplementary Figure 3:**
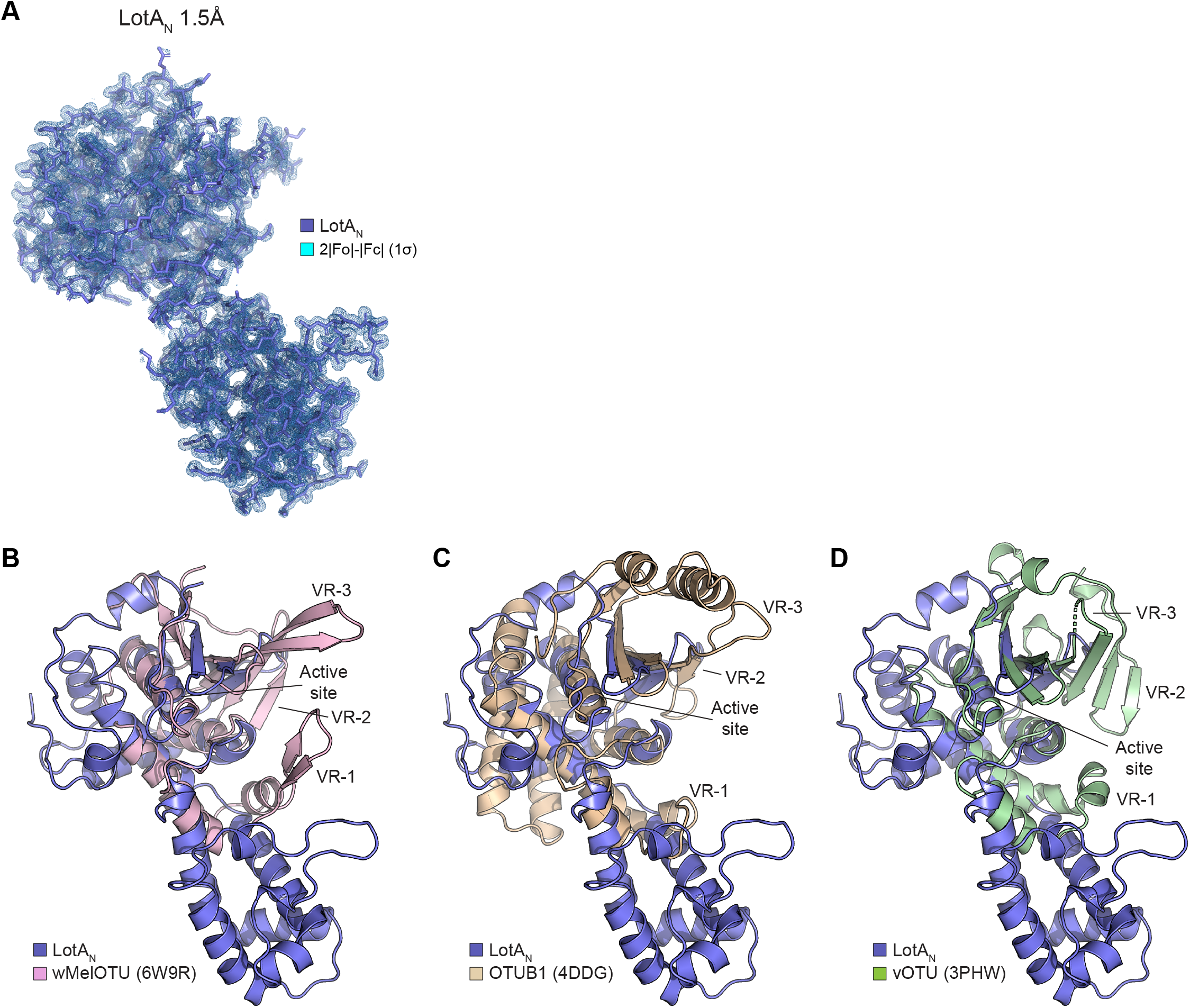
Crystal structure of the LotA_N_ OTU domain. A. 1.5Å crystal structure of LotA_N_ (1-294). The LotA_N_ model is shown in sticks with 2|Fo|-|Fc| electron density overlaid at 1α. B. Structural overlay of LotA_N_ (blue) with wMelOTU (pink) from *Wolbachia pipientis* wMel (PDB 6W9R). The aligned active sites are annotated and the unique variable regions of wMelOTU are indicated. C. Structural overlay of LotA_N_ (blue) with human OTUB1 (PDB 4DDG, tan). The aligned active sites are annotated and the unique variable regions of OTUB1 are indicated. D. Structural overlay of LotA_N_ (blue) with vOTU (green) from CrimeanCongo hemorrhagic fever virus (PDB 3PHW). The aligned active sites are annotated and the unique variable regions of vOTU are indicated.

**Supplementary Figure 4:**
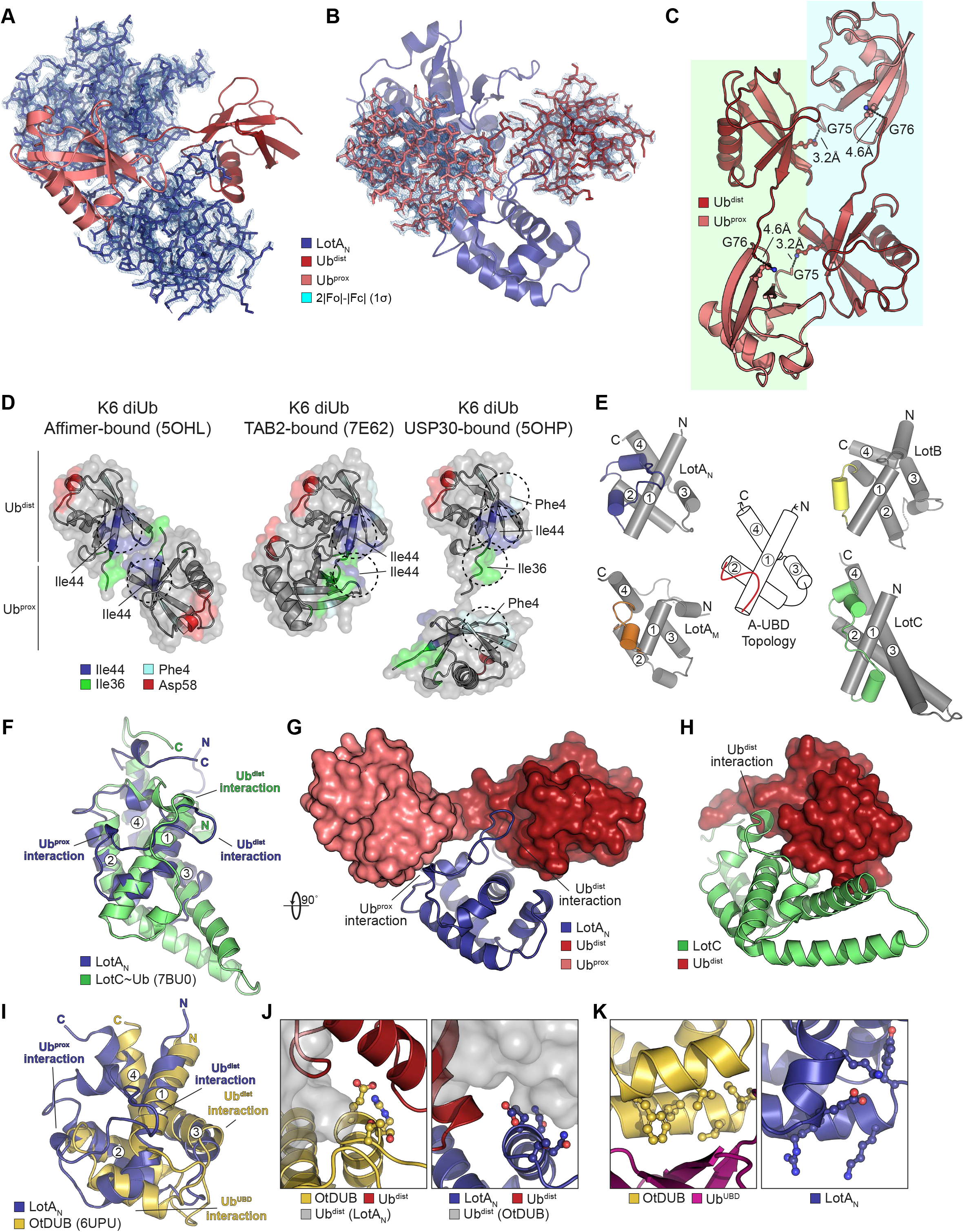
Crystal structure of the LotA_N_ OTU domain bound to K6 diUb. A. 2.8Å crystal structure of LotA_N_ (1-276) bound to K6 diUb. The LotA_N_ model (blue) is shown in sticks with 2|Fo|-|Fc| electron density overlaid at 1χρ, while the K6 diUb model (shades of red) is shown in cartoon. B. 2.8Å crystal structure of LotA_N_ (1-276) bound to K6 diUb. The K6 diUb model (shades of red) is shown in sticks with 2|Fo|-|Fc| electron density overlaid at 1χρ, while the LotA_N_ model (blue) is shown in cartoon. C. Cartoon representation of symmetry-related K6 diUb molecules. Distances between the Ub^dist^ C-terminus and Ub^prox^ K6 are shown within (blue or green) or across asymmetric units (blue to green). D. Crystal structures of K6 diUb bound to a K6-specific affimer (PDB 5OHL), TAB2 (PDB 7E62), or USP30 (PDB 5OHP), aligned by their distal Ub moieties (top) and shown side-by-side. Availability and orientation of common interaction surfaces are shown, with dashed circles indicating the surfaces utilized by protein interfaces. E. Underlying helical domain architecture of the Lot-class A-UBDs, with helices labeled and the Ub-binding α1-2 regions shown in color. F. Structural overlay of A-UBDs from LotA_N_ (blue) and LotC (green), with helices and Ub interaction surfaces labeled. G. Rotated view of the interaction between the LotA_N_ A-UBD (blue, cartoon) and K6 diUb (surface, shades of red), with Ub interaction surfaces labeled. H. Rotated view of the interaction between the LotC A-UBD (green, cartoon) and a Ub bound in the S1 site (surface, red), with Ub interaction surfaces labeled. I. Structural overlay of the A-UBDs from LotA_N_ (blue) and *Orientia tsutsugamushi* OtDUB (PDB 6UPU, gold), with helices and Ub interaction surfaces labeled. J. Side-by-side view of the OtDUB Ub^dist^ interaction surface on Helix 3 (gold) alongside the analogous, conserved region of LotA_N_ (blue). The bound Ub^dist^ molecules (red) would clash with additional Ub molecules (gray surface) overlaid from the opposite protein structure. K. Side-by-side view of the OtDUB Ub^UBD^ high affinity interaction site in the α1-2 region (gold) bound to Ub (maroon), alongside the distinct α1-2 region of LotA_N_ (blue).

**Supplementary Figure 5:**
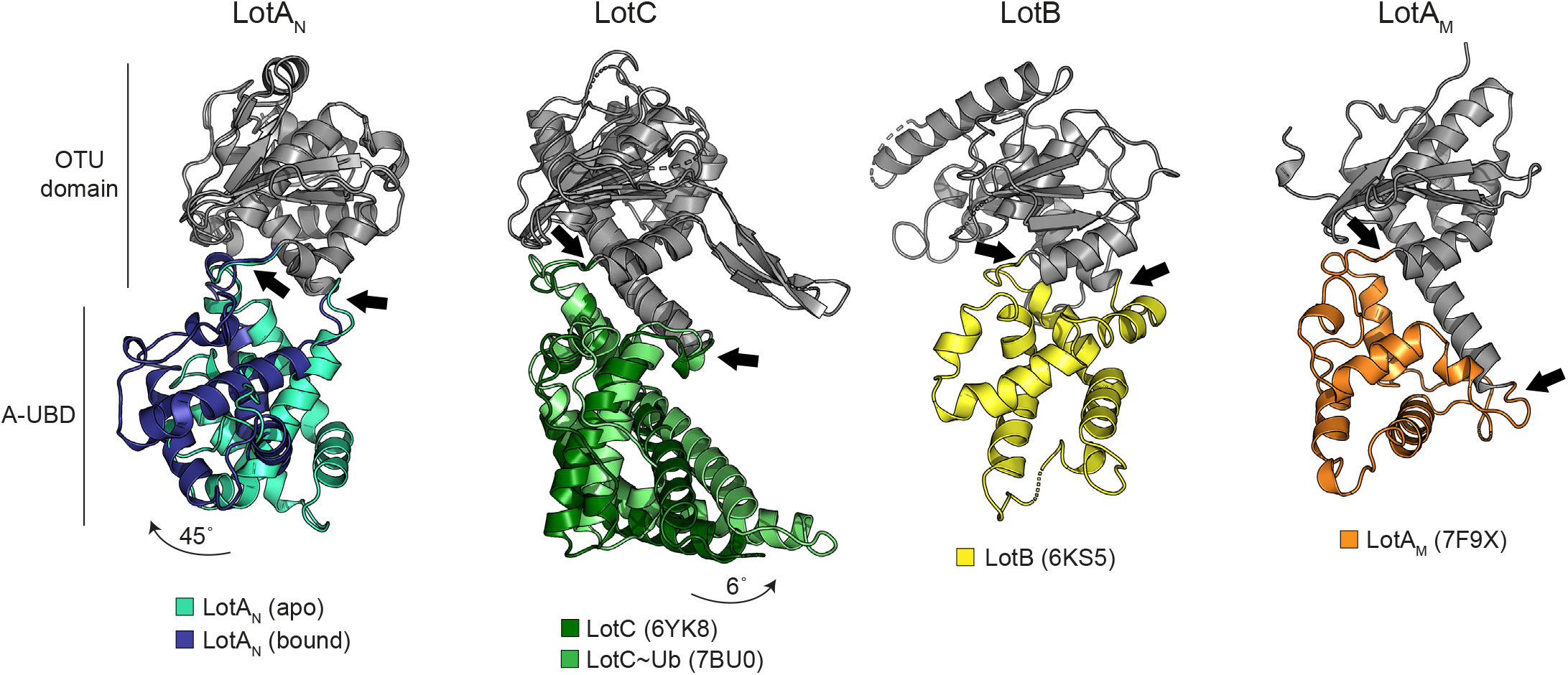
Conformational changes of LotA_N_ during catalysis. Side view of all structurally-characterized Lot-class DUBs highlighting hinge movement upon Ub binding for LotA_N_ (blue) and LotC (green), and putative hinge regions in the LotA_M_ (orange) and LotB (yellow) structures. Hinge regions between the OTU domain and A-UBD are indicated with black arrows.

**Supplementary Figure 6:**
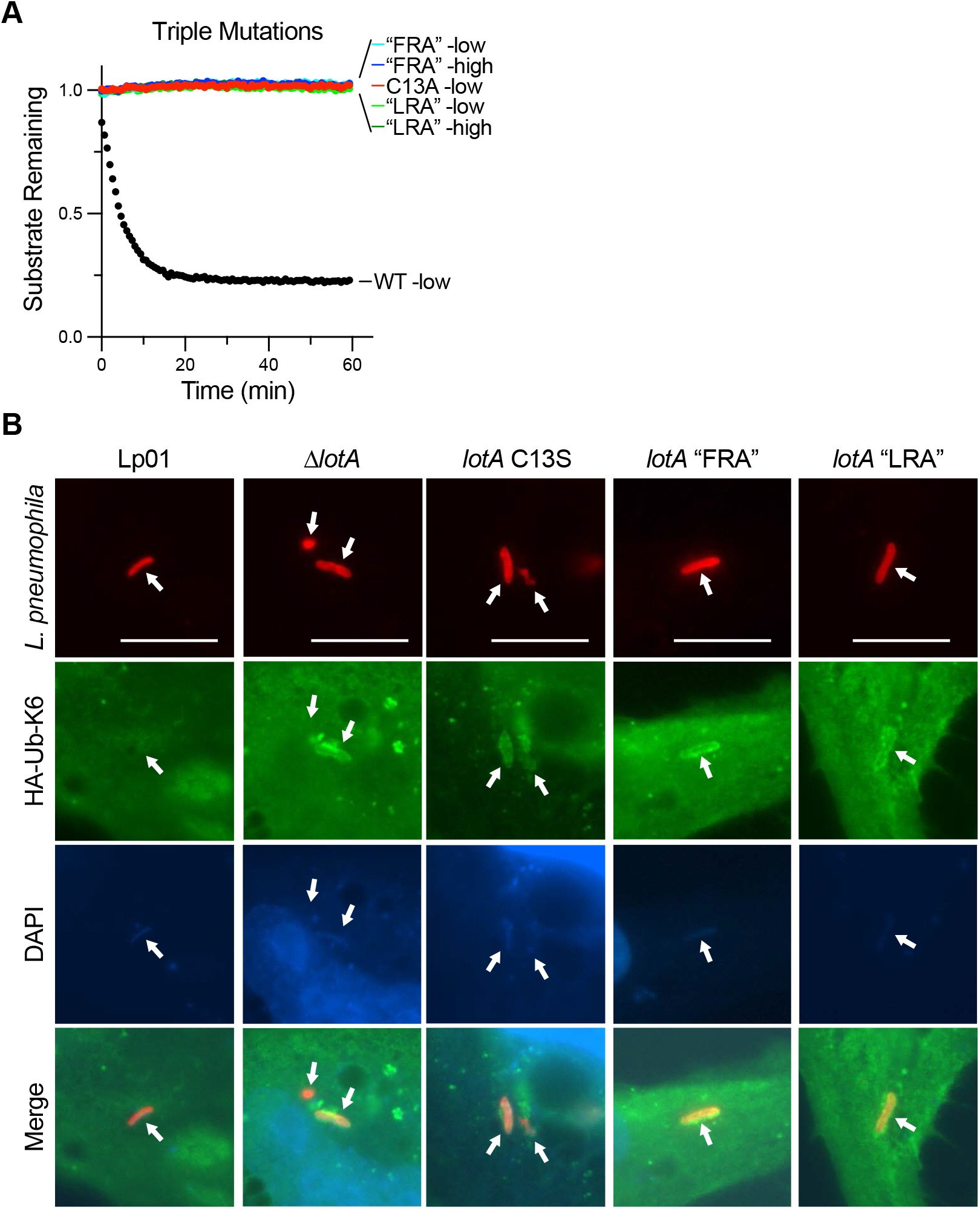
LotA_N_ restriction of K6 polyUb during *L. pneumophila* infection. A. Cleavage of K6 diUb by the indicated LotA_N_ variants at 10 nM (low) or 1 µM (high) concentration monitored by fluorescence polarization. The “FRA” triple mutant combines the LotA_N_ S1’-site F120A, S1’-site R145L, and hinge A193P mutations. The “LRA” triple mutant combines the LotA_N_ S1-site L128R, S1’-site R145L, and hinge A193P mutations. These data were collected in parallel with those presented in Fig. 3C, and the WT dataset is shown again for reference. B. Representative images of HeLa FcγRII cells infected with the indicated *L. pneumophila* strains at an MOI of 2 for 4 h. Fixed cells were stained for *L. pneumophila* (red), HA-Ub-K6 (green), and DNA (blue). Arrows indicate the position of a bacterium in each channel. Scale bars correspond to 10 µm.

**Table S1:** Data collection and refinement statistics.

|  | LotA<sub>N</sub> (1-294) | LotA<sub>N</sub> (1-276) : K6 diUb |
| --- | --- | --- |
| **Data collection** |  |  |
| Wavelength | 0.979460 | 0.979460 |
| Space group | *P* 4<sub>1</sub> 2<sub>1</sub> 2 | *P* 4 2 2 |
| Cell dimensions |  |  |
| *a*, *b*, *c* (Å) | 86.25, 86.25, 92.30 | 127.28, 127.28, 77.99 |
| $\alpha$, $\beta$, $\gamma$ (°) | 90, 90, 90 | 90, 90, 90 |
| Resolution (Å) | 39.07-1.50 (1.50-1.53) | 38.99-2.80 (2.80-2.95) |
| *R*<sub>merge</sub> | 0.035 (0.819) | 0.257 (1.15) |
| *I*/σ*I* | 26.6 (2.2) | 6.4 (2.0) |
| Completeness (%) | 99.9 (99.7) | 99.9 (100) |
| Redundancy | 6.5 (6.5) | 6.5 (6.7) |
| **Phasing** |  |  |
| Method | SAD | MR |
| SAD Resolution | 1.50 | - |
| SAD Anom completeness | 99.6 (99.5) | - |
| SAD Anom multiplicity | 3.4 (3.3) | - |
| SAD FOM | 0.733 | - |
| MR Search Models | - | LotA<sub>N</sub>, Ub (1UBQ) |
| **Refinement** |  |  |
| Resolution (Å) | 35.59-1.50 | 38.99-2.80 |
| No. unique reflections / test set | 56168 / 2895 | 16316 / 822 |
| *R*<sub>work</sub>/*R*<sub>free</sub> | 0.195/0.221 | 0.207/0.259 |
| No. atoms |  |  |
| Protein | 2174 | 3399 |
| Ligand/ion | 19 | 0 |
| Water | 286 | 59 |
| *B*-factors |  |  |
| Protein | 32.81 | 47.04 |
| Ligand/ion | 54.93 | - |
| Water | 39.12 | 33.63 |
| R.m.s. deviations |  |  |
| Bond lengths (Å) | 0.006 | 0.006 |
| Bond angles (°) | 0.77 | 1.04 |
| Ramachandran statistics |  |  |
| Favored / allowed / outliers | 96.7 / 3.3 / 0 | 97.2 / 2.8 / 0 |

Values in parentheses are for highest resolution shell.

**Table S2:**
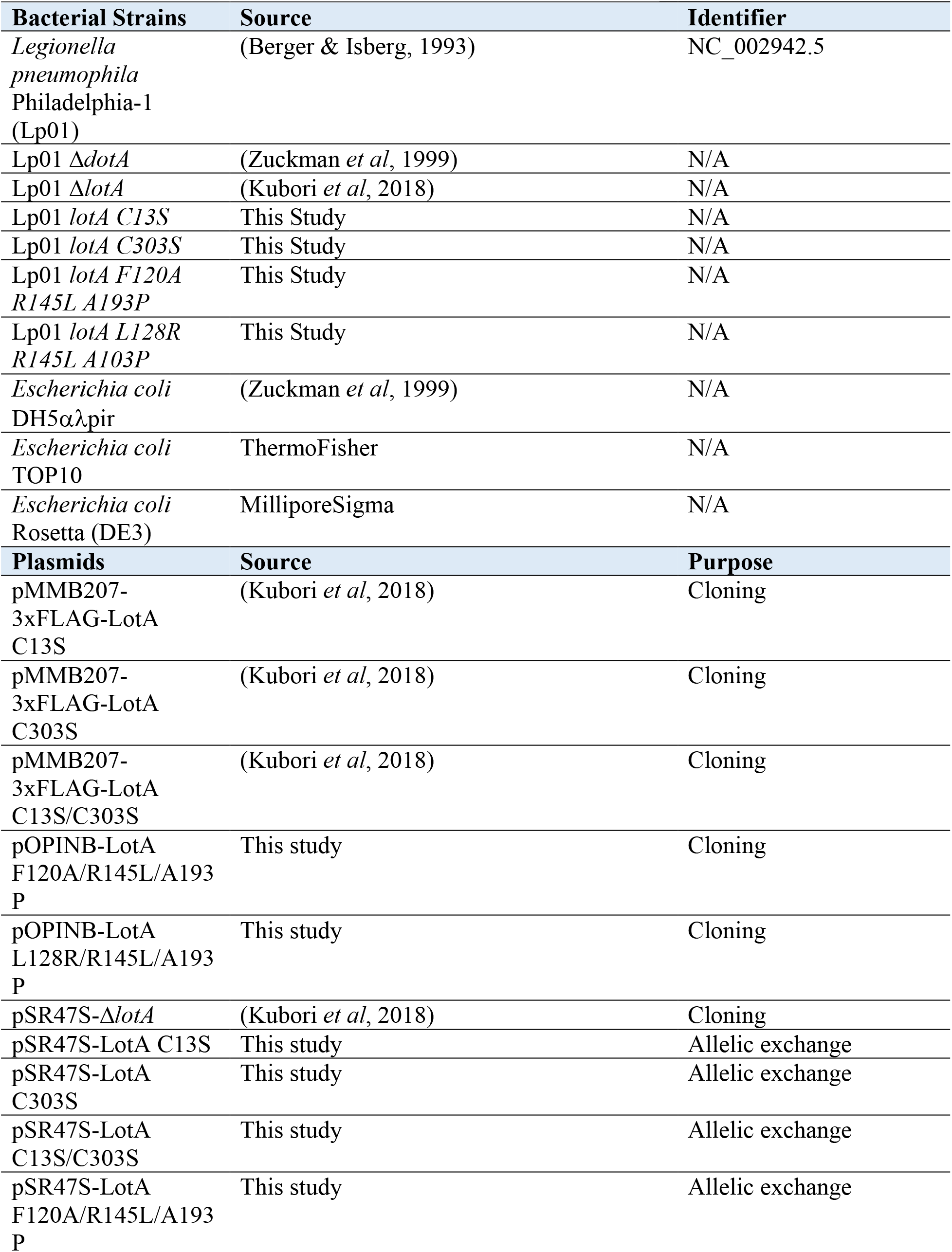
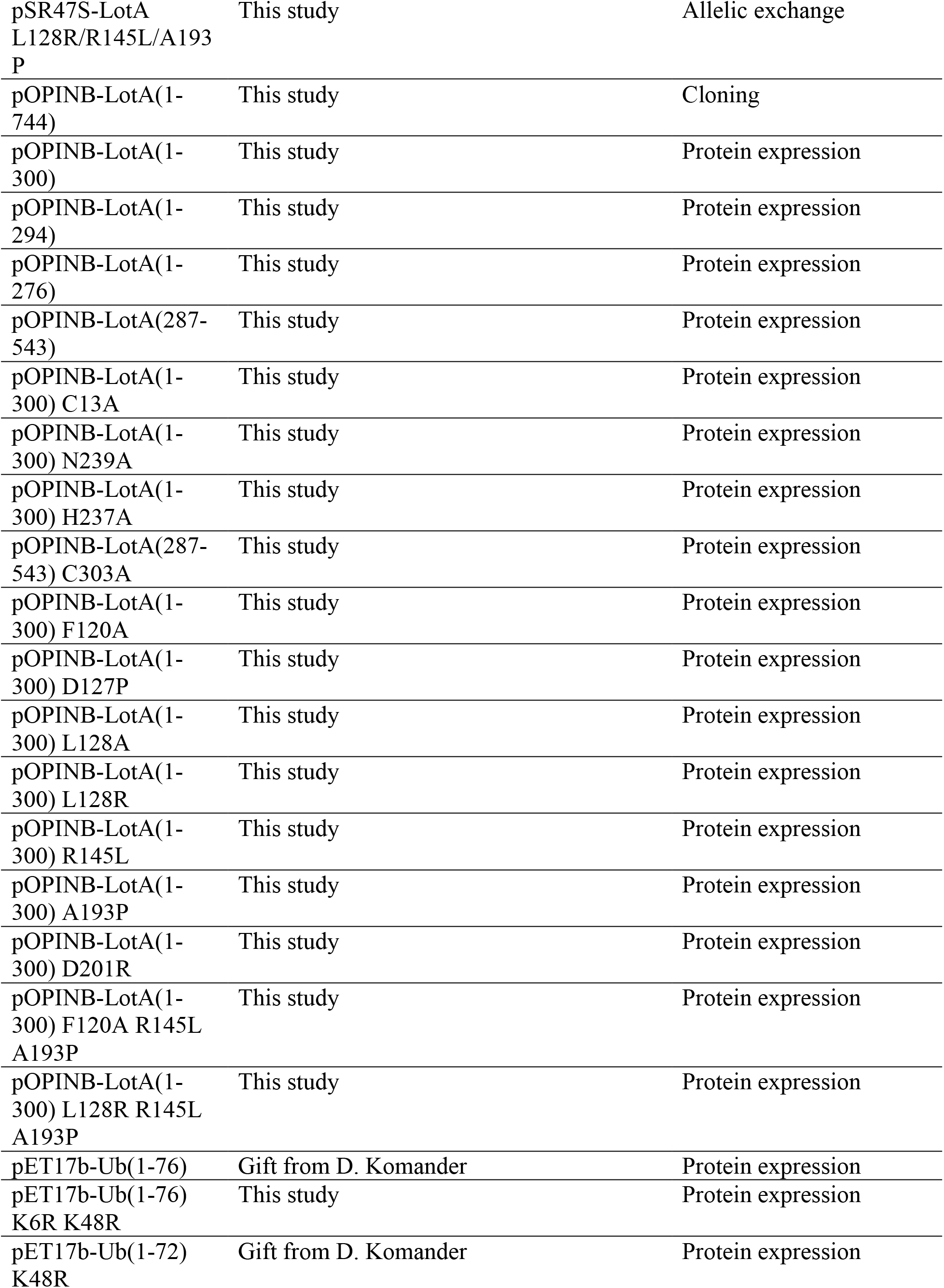
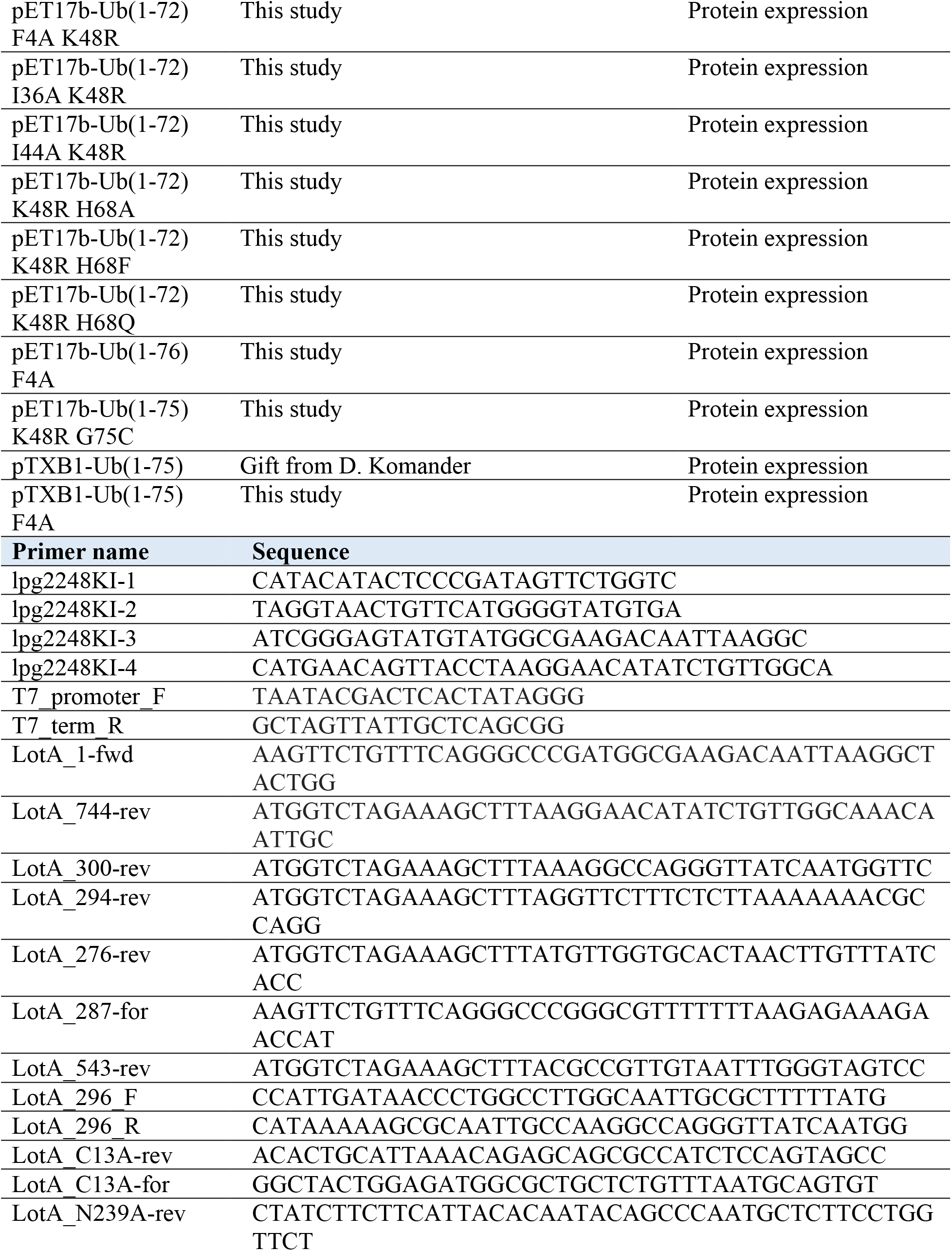
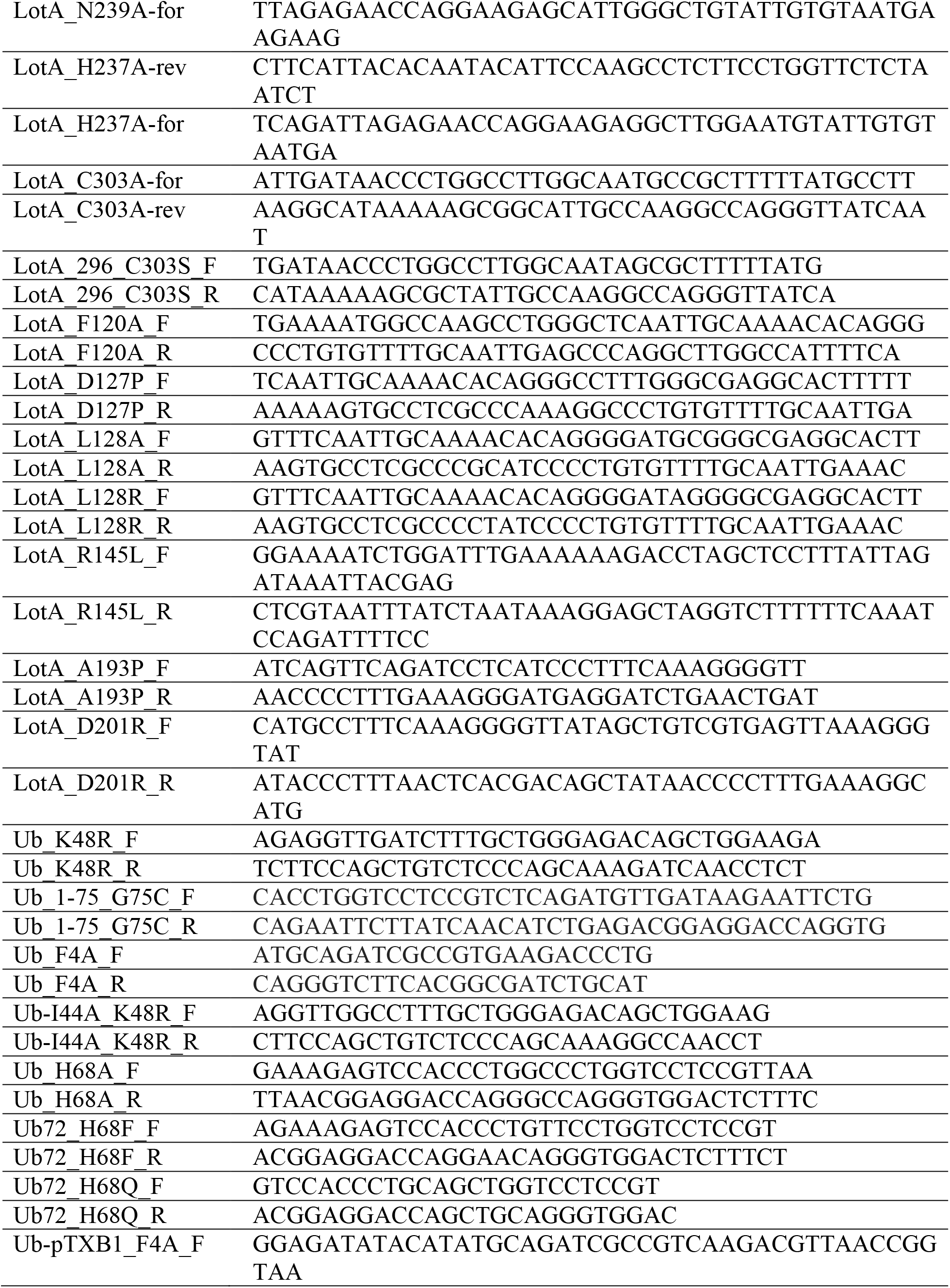
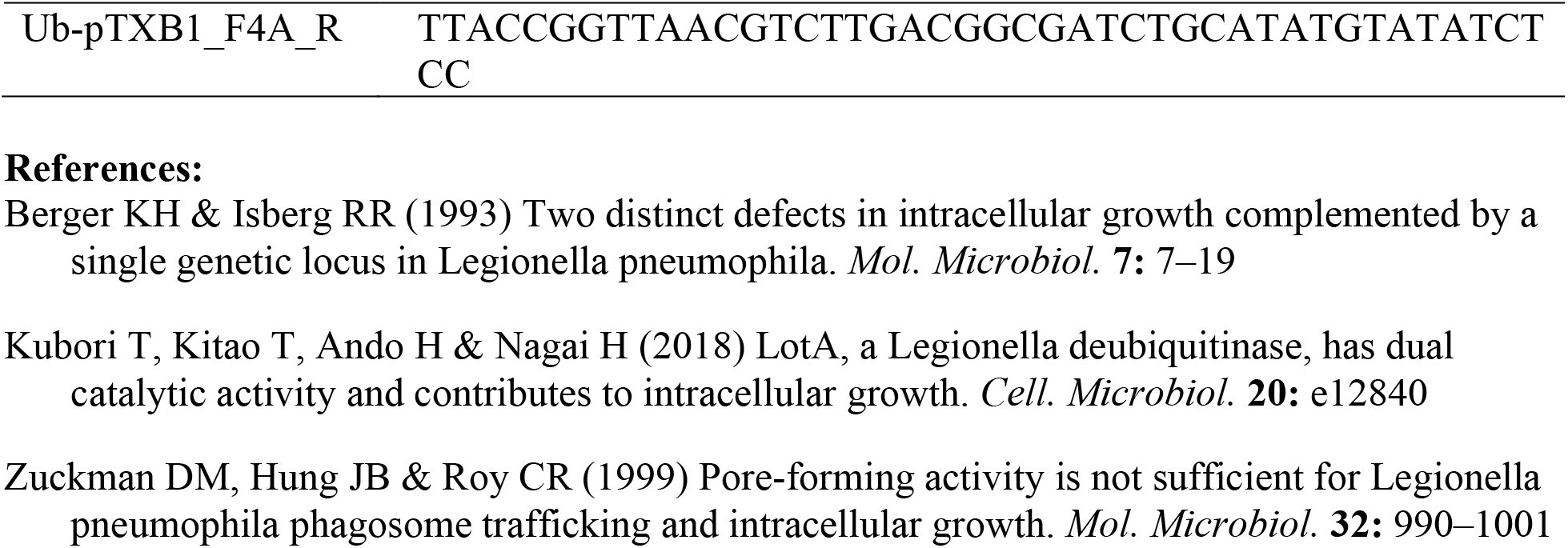
Bacterial strains and plasmids used in this study.

